# Unimeth: a unified transformer framework for accurate DNA methylation detection from nanopore reads

**DOI:** 10.64898/2025.12.05.692231

**Authors:** Shaokai Wang, Peng Ni, Yifu Xiao, Tao Sheng, Neng Huang, Yi Shu, Hong-Dong Li, Lu Shen, Xiao-Xin Yan, Jixian Zhai, Feng Luo, Jianxin Wang

**Affiliations:** School of Computer Science and Engineering, Central South University, Changsha 410083, China; Xiangjiang Laboratory, Changsha 410205, China; Hunan Provincial Key Lab on Bioinformatics, Central South University, Changsha 410083, China; Department of Data Sciences, Dana-Farber Cancer Institute, Boston, MA 02215, USA; Department of Biomedical Informatics, Harvard Medical School, Boston, MA 02115, USA; Department of Biology, School of Life Sciences, Southern University of Science and Technology, Shenzhen, China; Department of Neurology, Xiangya Hospital, Central South University, Changsha, China; Department of Anatomy and Neurobiology, Xiangya School of Basic Medical Sciences,Central South University, Changsha, China; School of Computing, Clemson University, Clemson, SC, 29634-0974, USA

**Author notes:** To whom correspondence should be addressed: Jianxin Wang. These authors contributed equally (Shaokai Wang, Peng Ni).

**Keywords:** nanopore sequencing, DNA methylation, methylation calling

## Abstract

Nanopore sequencing enables direct detection of DNA modifications from native DNA. However, accurate methylation calling across species, sequence contexts, modification types and chemistries remains challenging. We developed Unimeth, a transformer-based framework that jointly processes raw signals and basecalled sequences to predict methylation at all target sites within read patches. Unimeth uses multi-phase training that combines signal pre-training, methylation fine-tuning and site-level calibration. We evaluated Unimeth for 5mC and 6mA detection using public and in-house datasets spanning 14 organisms, R9.4.1 and R10.4.1 nanopore chemistries, and wild-type, mutant and enzyme-treated samples. Unimeth improved plant 5mC detection in non-CpG contexts, reduced false-positive calls in low-methylation samples and maintained high performance for 5mCpG detection. We also sequenced an Alzheimer’s disease brain sample and further validated Unimeth. For 6mA, Unimeth reduced background calls while retaining signals for nucleosome profiling and gene-level analysis. Unimeth provides an open-source framework for methylation detection across modification types, species and nanopore chemistries.

## Introduction

DNA methylation, including 5-methylcytosine (5mC) and N^6^-methyladenine (6mA), is involved in diverse biological processes in bacteria, mammals, and plants^1–3^. Traditionally, DNA methylation has been extensively studied using microarray or short-read sequencing methods that rely on specialized library preparation, such as chemical conversion, antibody enrichment, or enzyme digestion^4,5^. However, these preparation steps can introduce biases in methylation profiling^5^. For example, methylated DNA immunoprecipitation-sequencing (MeDIP-seq), which enriches DNA fragments using antibodies against 5mC, lacks single-nucleotide resolution and is biased toward highly methylated regions^6^. Bisulfite sequencing (BS-seq), although widely used for cytosine methylation profiling, can suffer from incomplete conversion and DNA degradation, leading to uneven coverage and false-positive calls^4,5,7,8^.

Long-read sequencing technologies, including Oxford Nanopore Technologies (ONT) sequencing and PacBio single-molecule real-time (SMRT) sequencing, have emerged as powerful platforms for DNA methylation detection^9,10^. Compared with short-read approaches, long-read sequencing can analyze native DNA molecules directly, avoiding chemical conversion or enrichment steps and reducing preparation-related biases^5,11^. It also supports simultaneous detection of multiple methylation types at single-molecule and single-nucleotide resolution. In addition, long reads facilitate epigenetic profiling across repetitive regions, including transposons^12^ and centromeres^13,14^, and across megabase-scale haplotype blocks^15,16^.

A range of computational tools has been developed to detect DNA methylation from nanopore reads^17–20^. These tools include statistical approaches that compare native and unmodified DNA signals, as well as machine-learning and deep-learning methods that classify methylation states at target sites^21–23^. With the release of ONT R10.4.1 chemistry, several tools have been developed or updated for methylation calling, including the official ONT tool Dorado, DeepMod2^24^, Rockfish^25^ and DeepPlant^26^. These methods have achieved high accuracy in selected datasets^19,20^, and recent benchmarks have shown that their relative performance varies among modification types and datasets^18,27^.

Despite these advances, current nanopore methylation callers have generally been developed for specific species, sequence contexts, modification types or sequencing chemistries. Dorado supports multiple DNA modifications, including 5mC, 4mC, 6mA and 5-hydroxymethylcytosine (5hmC), but its performance varies across modification types and sequence contexts^27^. Rockfish^25^ and DeepMod2^24^ perform well for 5mCpG detection in human and mouse data but are restricted to CpG sites. They therefore do not address CHG and CHH methylation (H = A, C or T), which has important roles in transposon silencing and gene regulation in plants^28^. Detecting non-CpG methylation is particularly challenging because methylated cytosines often represent only a small fraction of observations in these contexts, especially at CHH sites, leading to strong class imbalance. Several methods have attempted to address this issue by modifying the training data. DeepSignal-plant^29^ uses semi-supervised learning to generate pseudo-labeled samples and rebalance training sets, whereas DeepPlant^26^ incorporates data from multiple species and uses higher methylation-frequency cutoffs to select confident training sites. In addition, 6mA detection remains constrained by species and chemistry specificity. For example, mCaller was developed for bacterial 6mA detection using R9 reads^30^.

We therefore developed Unimeth, a unified transformer-based framework for DNA methylation detection from nanopore reads. First, Unimeth uses a patch-based approach that jointly processes raw signals and basecalled sequences across continuous read segments. This approach enables simultaneous prediction of all targeted methylation sites within each patch and allows the model to use neighbouring sequence and signal context. Second, Unimeth uses a multi-phase training strategy. During pre-training, the model learns signal features through a basecalling-like task. During fine-tuning, high-confidence methylated and unmethylated sites are used to train methylation prediction. When site-level methylation measurements are available, a calibration stage further allows sites with intermediate methylation levels to contribute to additional fine-tuning.

We trained and evaluated Unimeth models for 5mC and 6mA detection using public and in-house datasets from 14 organisms, including 10 plants, 2 mammals, 1 heterolobosean protist and 1 bacterium. These datasets included R10.4.1 5kHz, R10.4.1 4kHz and R9.4.1 nanopore reads from wild-type, mutant and enzyme-treated samples. Across the plant datasets, Unimeth improved non-CpG 5mC detection and reduced false-positive calls in low-methylation samples. It maintained high 5mCpG accuracy in human and mouse datasets. We also sequenced an Alzheimer’s disease brain sample and validated CpG methylation calls from Unimeth against matched BS-seq. For 6mA, Unimeth reduced background calls across diverse datasets and preserved signals used to profile nucleosome organization and methylation patterns across genes. We provide Unimeth as an open-source tool with defined data partitions, training and inference code, pretrained models and support for standard nanopore input and output formats.

## Results

### The framework of Unimeth

Unimeth uses a patch-based strategy to detect DNA methylation from nanopore reads (Fig. 1a). Nanopore reads are divided into overlapping patches, each comprising a basecalled sequence segment and its corresponding raw-signal segment. Context tokens are inserted at target bases, including CpG, CHG and CHH cytosines for 5mC detection and adenines for 6mA detection. A transformer encoder processes normalized signal embeddings, whereas a transformer decoder integrates signal, sequence and context information to classify each marked site as methylated or unmethylated. This design enables multiple sites to be called together while retaining their local sequence and signal context.

**Fig. 1.**
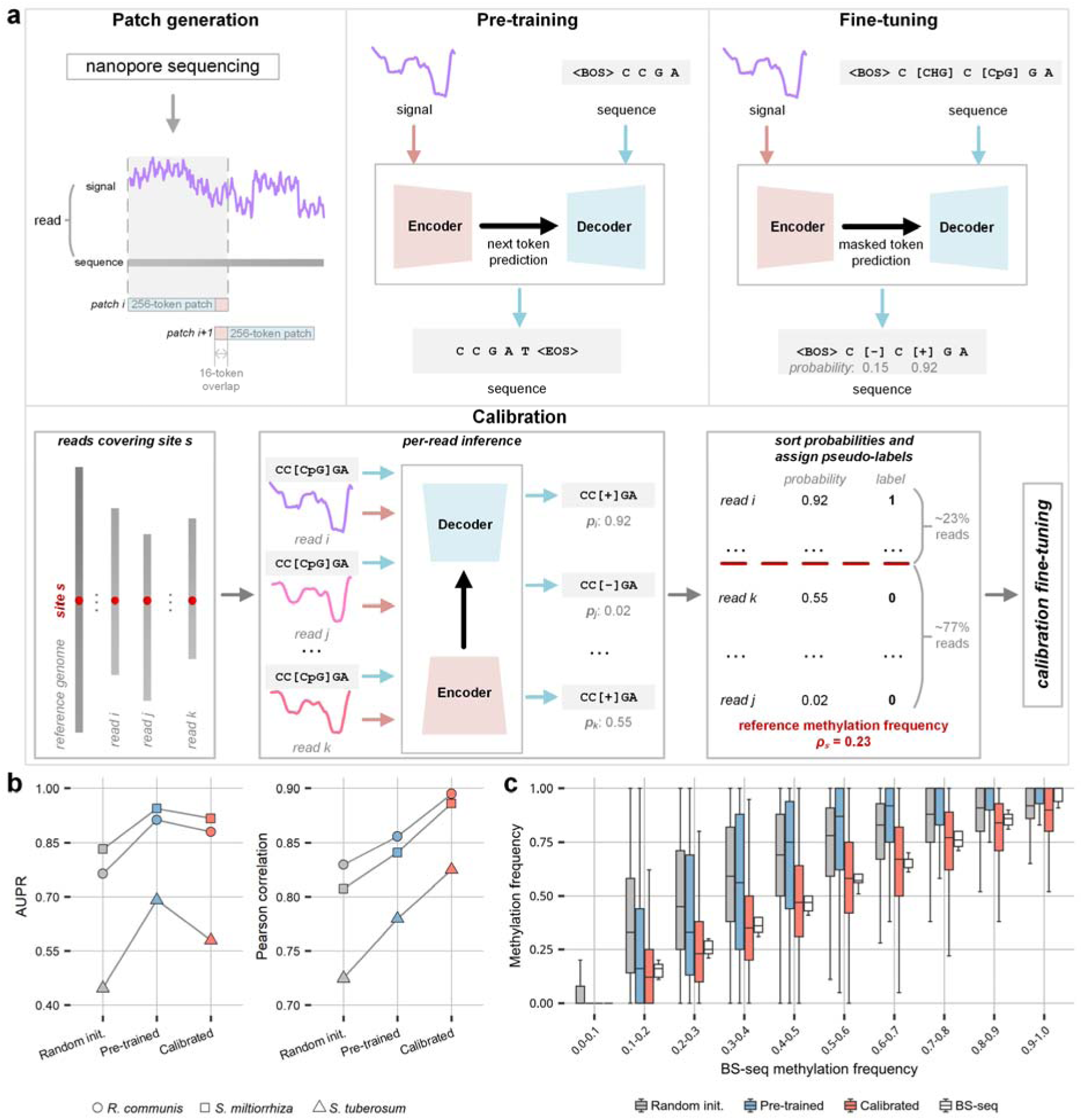
The framework and multi-phase training strategy of Unimeth. **a** Patch-based model architecture and training phases. Nanopore reads are divided into overlapping patches of 256 tokens, each paired with the corresponding signal segment. During pre-training, signal and sequence patches are used for next-token reconstruction. During fine-tuning, context tokens mark target sites ([CpG], [CHG] and [CHH] for 5mC, and [A] for 6mA), and the model predicts methylated ([+]) or unmethylated ([-]) labels. When site-level references are available, calibration sorts read-level methylation probabilities within each genomic site and assigns pseudo-labels according to the reference methylation frequency for additional fine-tuning. **b** Effects of pre-training and calibration on CHH methylation calling in *R. communis*, *S. miltiorrhiza* and *S. tuberosum*. Random init. and Pre-trained used the same fine-tuning data and procedure but started from random and pre-trained weights, respectively; Calibrated included an additional calibration step. Read-level AUPR (left) and site-level Pearson correlation with BS-seq at 20× nanopore coverage (right) are shown. Site-level values represent the mean of five subsampling replicates. **c** Site-level CHH methylation frequencies predicted by the three model variants and measured by BS-seq in *S. miltiorrhiza*, stratified by BS-seq methylation frequency. Boxes show interquartile ranges, centre lines indicate medians, and whiskers extend to values within 1.5 times the interquartile range; outliers are not shown.

Unimeth uses up to three training phases (Fig. 1a). Pre-training reconstructs nucleotide sequences from raw signal embeddings, providing an initialization that captures relationships between nanopore signals and nucleotide sequences. During fine-tuning, context tokens are inserted at candidate methylation sites, and the model learns to predict methylated or unmethylated labels from high-confidence training sites. When quantitative site-level references such as BS-seq are available, calibration generates pseudo-labels from read-level methylation probabilities and the corresponding reference methylation frequencies. This allows sites with intermediate methylation levels to contribute to additional fine-tuning. Once trained, Unimeth accepts POD5 or SLOW5 signal files together with basecalled BAM files and outputs read-level modification calls in modBAM format and site-level methylation frequencies in BED format (Supplementary Fig. 1).

We examined the effects of pre-training and calibration using held-out chromosomes or contigs from the R10.4.1 5kHz DeepPlant datasets for *R. communis*, *S. miltiorrhiza* and *S. tuberosum*^26^ (Supplementary Data 1 and 2). All model variants used the same high-confidence training sites and the same initial fine-tuning procedure (Methods). Pre-training generally improved read-level area under the precision-recall curve (AUPR) and area under the receiver operating characteristic curve (AUC) across methylation contexts, with the clearest improvements for CHH (Fig. 1b and Supplementary Fig. 2). Calibration reduced false-positive calls and increased read-level precision, although sensitivity was lower and AUPR did not improve (Fig. 1b and Supplementary Fig. 2). At the site level, calibration consistently increased Pearson correlation and generally reduced root mean square error (RMSE) across species and methylation contexts, with the clearest improvements for CHH (Fig. 1b and Supplementary Fig. 3). Reducing false-positive calls may help correct methylation overestimation at intermediately methylated sites, bringing predictions closer to the corresponding BS-seq measurements (Fig. 1c and Supplementary Fig. 4).

### Read-level evaluation of 5mC detection in plants

We evaluated read-level 5mC detection across all nine R10.4.1 5kHz plant datasets from DeepPlant^26^, which included species with different methylation levels, particularly in CHH contexts (Supplementary Data 3). The three training species, *R. communis*, *S. miltiorrhiza* and *S. tuberosum*, were evaluated on held-out chromosomes or contigs, whereas the remaining six species were used to assess cross-species generalization (Supplementary Data 2). We compared Unimeth with Dorado and DeepPlant across CpG, CHG and CHH contexts, and with DeepMod2 and Rockfish for CpG sites (Methods).

Across the nine plant datasets, Unimeth showed consistently high read-level performance (Fig. 2 and Supplementary Fig. 5). For CpG and CHG detection, Unimeth achieved the highest AUC in eight of nine species, with mean AUCs of 0.9862 and 0.9910, respectively. For CHH detection, Unimeth achieved the highest AUC and AUPR in all nine species, with mean values of 0.9817 and 0.5049, compared with 0.9254 and 0.2709 for DeepPlant and 0.9012 and 0.1336 for Dorado. Among the six species not used for training, Unimeth also showed high AUCs across CpG, CHG and CHH contexts. For CHH sites in these species, Unimeth achieved a mean AUPR of 0.3614, compared with 0.1120 for DeepPlant and 0.0401 for Dorado. Additional read-level metrics, including accuracy, specificity, precision and sensitivity, were consistent with these comparisons (Supplementary Fig. 5). In independent plant datasets generated in-house, Unimeth achieved the highest mean AUC in each of the CpG, CHG and CHH contexts, with the clearest improvements in CHG and CHH detection (Methods and Supplementary Fig. 6). To evaluate an older nanopore chemistry, we trained an R9.4.1 model and tested it on *A. thaliana* and *O. sativa* reads (Supplementary Data 2). In this evaluation, Unimeth outperformed DeepSignal-plant across all three contexts, with the largest difference in mean CHH AUPR (0.7215 versus 0.2257; Supplementary Fig. 7).

**Fig. 2.**
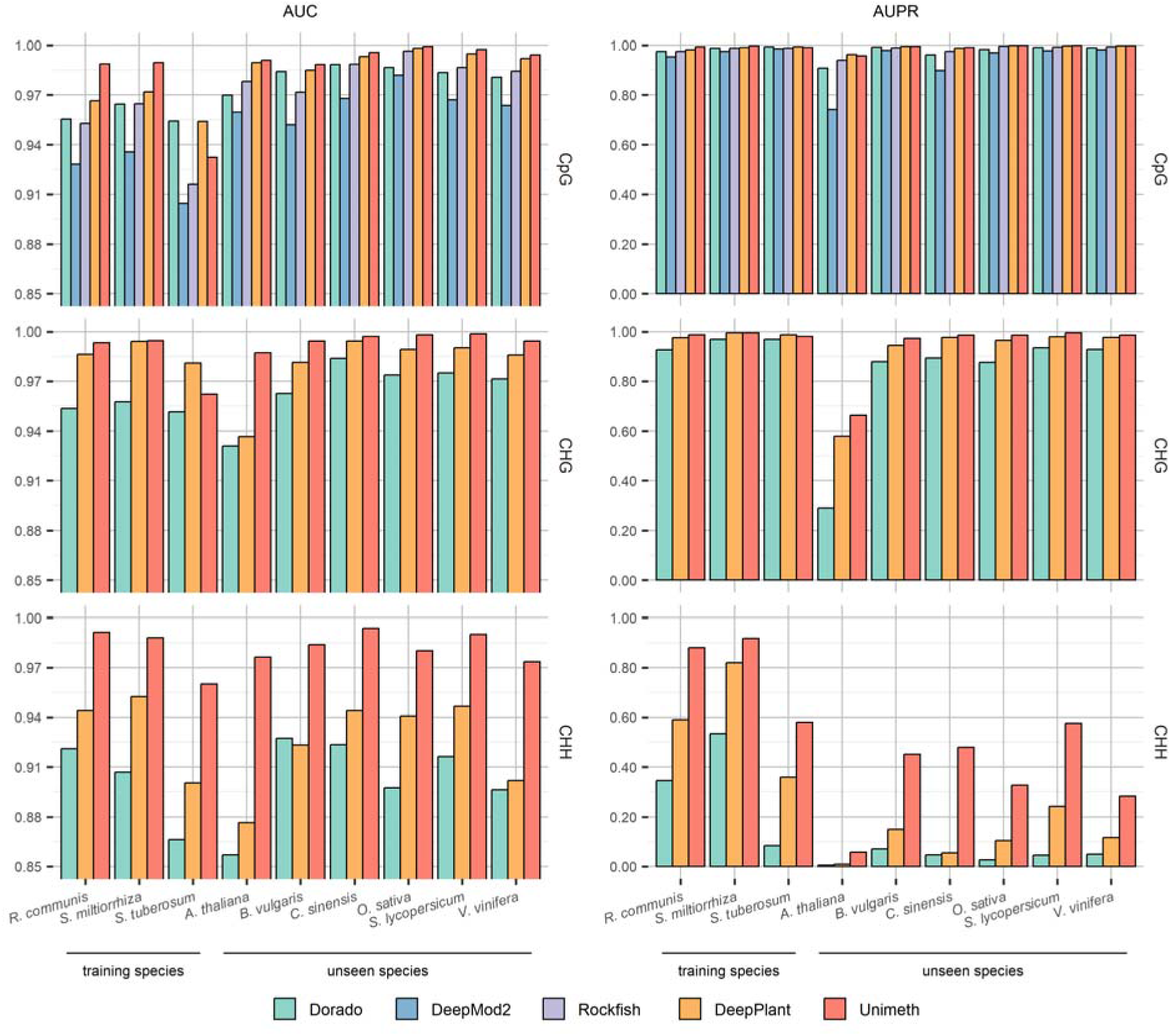
Read-level evaluation of 5mC detection in plants using ONT R10.4.1 5kHz reads. AUC and AUPR are shown for CpG, CHG and CHH methylation detection across the nine DeepPlant datasets. The first three species were used for model training but evaluated on held-out chromosomes or contigs, whereas the remaining six species were not used for training.

### Site-level evaluation of 5mC detection in plants

We evaluated site-level 5mC quantification by aggregating read-level calls at cytosine sites and comparing methylation frequencies with matched BS-seq data (Methods). In the nine R10.4.1 5kHz plant datasets from DeepPlant^26^, Unimeth generally showed the highest Pearson correlations across nanopore coverages, especially for CHG and CHH methylation at lower coverage (Fig. 3a and Supplementary Data 4). Using all available nanopore reads, the mean Pearson correlations were 0.9605, 0.9618 and 0.8574 for CpG, CHG and CHH, respectively, compared with 0.9546, 0.9283 and 0.6711 for Dorado and 0.9482, 0.9300 and 0.8298 for DeepPlant. Spearman correlation, coefficient of determination and root mean square error (RMSE) showed the same overall pattern (Supplementary Data 4).

**Fig. 3.**
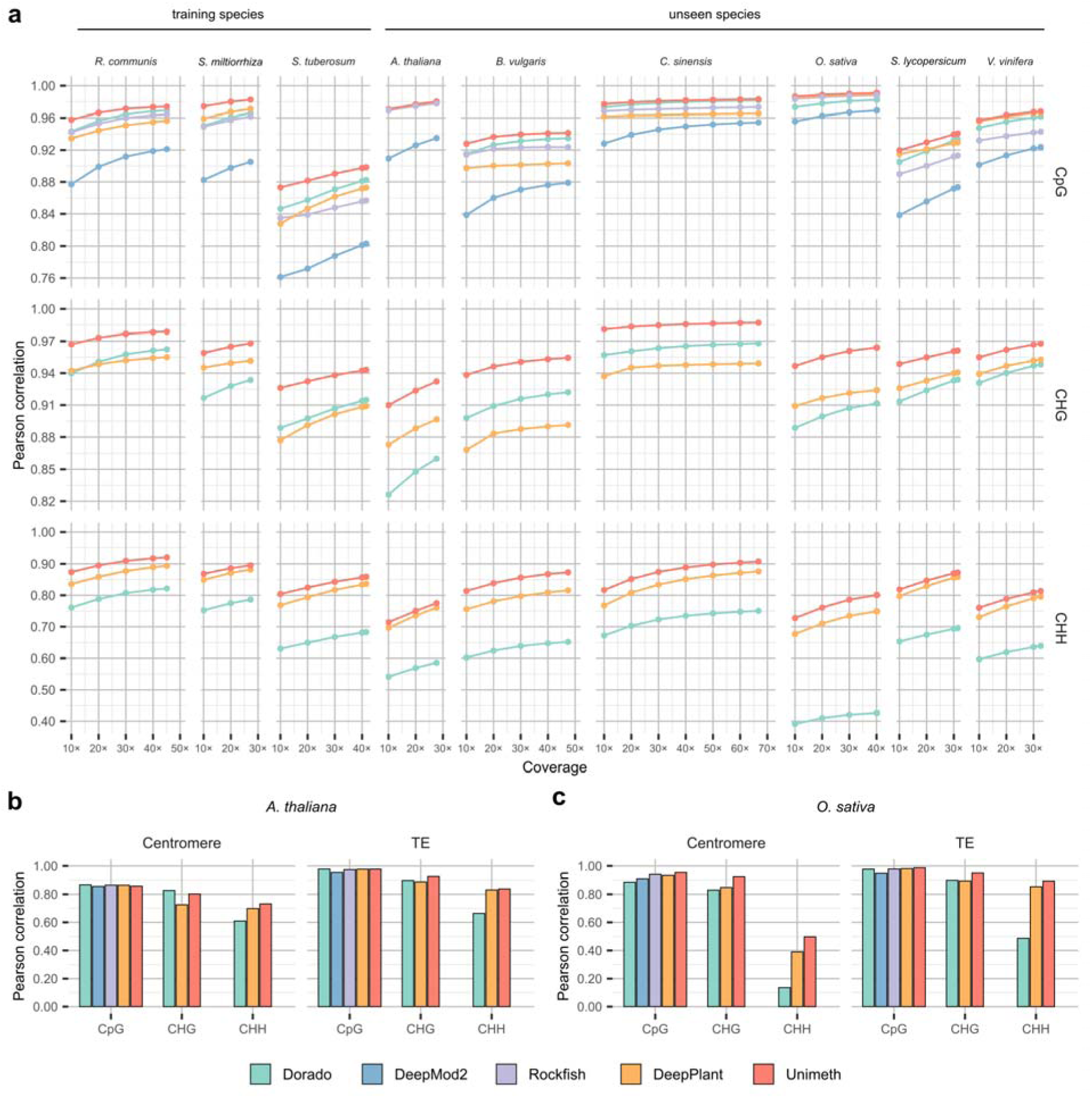
Site-level evaluation of Unimeth and other methods for plant 5mC detection using nanopore R10.4.1 5kHz reads. **a** Pearson correlations between methylation frequencies estimated from nanopore reads and matched BS-seq data at cytosine sites in CpG, CHG and CHH contexts across nanopore read coverages. The first three species were used for model training but evaluated on held-out chromosomes or contigs, whereas the remaining six species were not used for training. Values at subsampled coverage levels represent the mean of five subsampling replicates; standard deviations are provided in Supplementary Data 4. **b,c** Pearson correlations between methylation frequencies estimated from nanopore reads and matched BS-seq data in repetitive genomic regions of *A. thaliana* (**b**) and *O. sativa* (**c**), including centromeric and transposable element (TE) regions. Analyses were performed using 80× coverage of in-house nanopore reads. Values represent the mean of five subsampling replicates.

In independent in-house R10.4.1 5kHz datasets from *A. thaliana*, *O. sativa* and *Musa* spp., Unimeth reached mean Pearson correlations of 0.9855, 0.9618 and 0.8435 for CpG, CHG and CHH contexts, respectively, at 80× coverage (Supplementary Fig. 8 and Supplementary Data 5). The next-highest values were 0.9832 for Dorado in CpG and 0.9281 and 0.8160 for DeepPlant in CHG and CHH. An R9.4.1 Unimeth model was also evaluated on *A. thaliana* and *O. sativa* reads. At 80× coverage, it outperformed DeepSignal-plant with mean Pearson correlations of 0.9920, 0.9773 and 0.9202 for CpG, CHG and CHH, respectively, compared with 0.9850, 0.9708 and 0.9036 for DeepSignal-plant (Supplementary Fig. 9 and Supplementary Data 6).

We also compared genomic cytosine coverage in nanopore and matched BS-seq data. In the public and in-house R10.4.1 5kHz plant datasets, the fraction of profiled cytosines increased with nanopore depth and generally approached or exceeded the matched BS-seq fraction across CpG, CHG and CHH contexts (Supplementary Figs. 10 and 11). In repetitive genomic regions of *A. thaliana* and *O. sativa*, including centromeric and transposable element regions, nanopore reads covered a larger fraction of cytosines than BS-seq (Supplementary Fig. 12), and Unimeth generally achieved the highest correlations and lowest RMSEs for site-level methylation frequency quantification (Fig. 3b,c and Supplementary Fig. 13). In *O. sativa*, Unimeth achieved the highest correlations and lowest RMSEs for CHG and CHH methylation in both transposable elements and centromeres (Supplementary Data 7).

### Unimeth reduces false-positive 5mC calls in plants

False-positive methylation calls can have a large effect in plant genomes, particularly in non-CpG contexts where methylated cytosines are often less abundant. We therefore evaluated read-level false-positive rates (FPRs) using high-confidence unmethylated cytosines from matched BS-seq data (Methods). In the nine public R10.4.1 5kHz plant datasets, Unimeth showed low FPRs across contexts, with the clearest reductions in CHG and CHH sites (Fig. 4a). For CHH sites, the mean FPR was 0.0075 for Unimeth, compared with 0.0142 for DeepPlant and 0.0563 for Dorado. CpG FPRs were low for several methods, and Rockfish or DeepPlant had lower values in some datasets. In-house R10.4.1 5kHz datasets and R9.4.1 plant reads showed the same overall pattern (Supplementary Figs. 14 and 15).

**Fig. 4.**
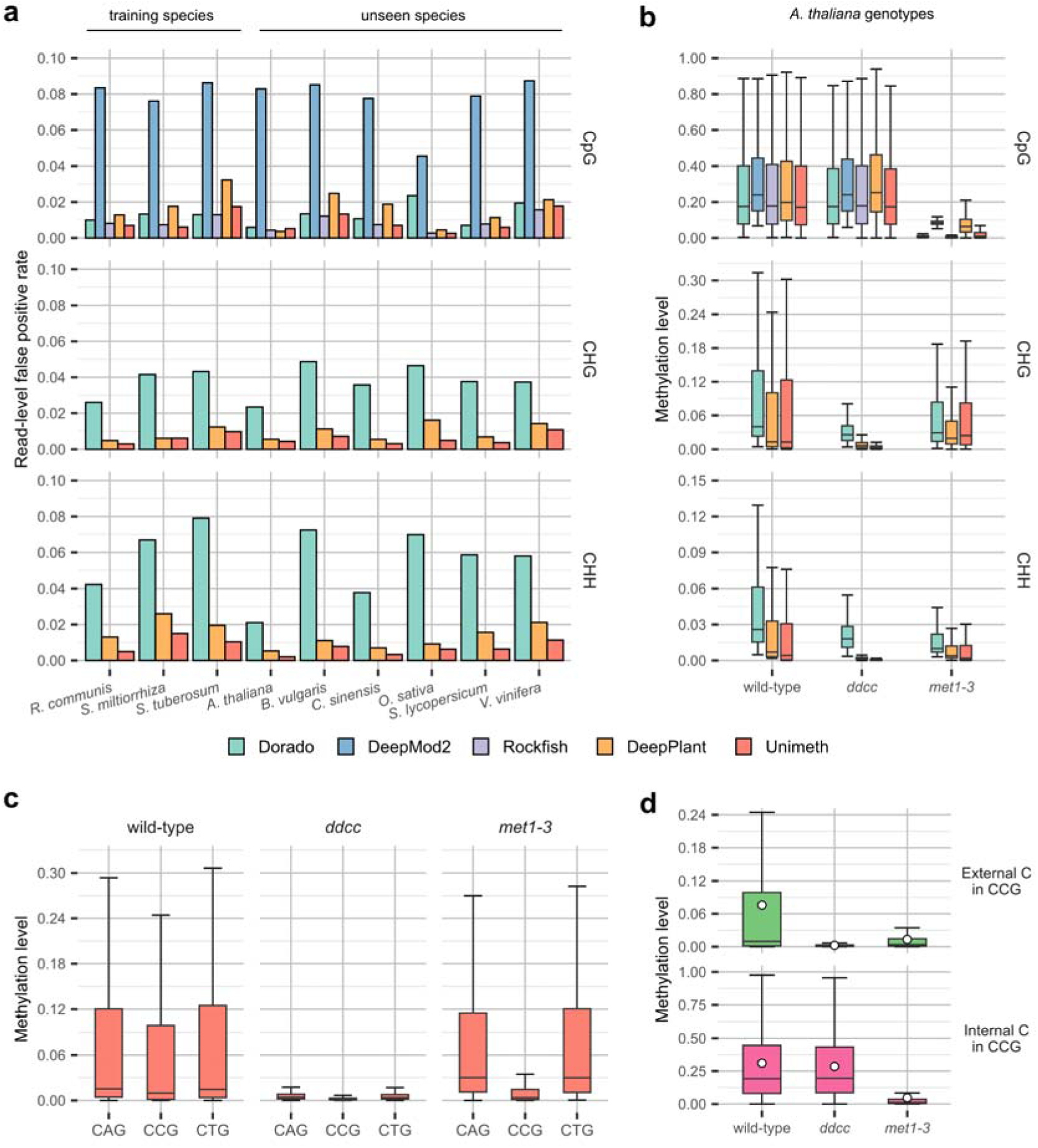
Unimeth reduces false-positive 5mC calls in plants. **a** Read-level false-positive rates of 5mC detection methods on plant nanopore R10.4.1 5kHz reads across CpG, CHG, and CHH contexts. **b** Methylation levels in *A. thaliana* Col-0 wild type and methylation-deficient mutants *ddcc* and *met1-3*. Methylation levels were calculated for 10-kb windows as the ratio of methylated calls to total covered calls within each window. **c** Unimeth-estimated methylation levels at CAG, CCG and CTG sites in the same wild-type and mutant samples, calculated for 10-kb genomic windows. **d** Unimeth-estimated window-level methylation distributions for paired cytosines in CCG motifs, comparing the external CHG-context cytosine with the internal CpG-context cytosine. White circles indicate count-weighted methylation levels across all covered calls. For **b-d**, boxes show interquartile ranges, centre lines show medians, and whiskers indicate 1.5 times the interquartile range.

Methylation-deficient *A. thaliana* mutants provided an additional test of background 5mC calling. In R10.4.1 5kHz data from Col-0 wild type, *drm1 drm2 cmt2 cmt3* (hereafter termed *ddcc*) and *met1-3* mutants, Unimeth recovered the expected loss of non-CpG methylation in *ddcc* and reduced CpG methylation in *met1-3*, while retaining higher methylation in the corresponding wild-type contexts (Fig. 4b). Residual methylation in mutant samples was lower with Unimeth than with the other methods, especially in CHG and CHH contexts. Using Unimeth calls, motif-level analyses further showed near depletion of CAG, CCG and CTG methylation in *ddcc* and selective reduction of CCG methylation in *met1-3* (Fig. 4c and Supplementary Fig. 16a)^31,32^. In CCG motifs, Unimeth separated the external CHG-context cytosine from the internal CpG-context cytosine, consistent with the dependence of external CCG methylation on internal CpG methylation (Fig. 4d and Supplementary Fig. 16b)^33^. Published R9.4.1 *A. thaliana* mutant data showed the same trend (Supplementary Fig. 17). These analyses support reduced background 5mC calling by Unimeth in low-methylation and methylation-depleted plant samples.

### Unimeth accurately profiles 5mCpG across species and human haplotypes

We evaluated CpG methylation calling in mammalian and bacterial genomes. For the human R10.4.1 5kHz evaluation, Unimeth was trained on HG002 data excluding chromosome 1, and tested on chromosome 1 from HG001-HG004 (Supplementary Data 2). Unimeth achieved the highest average read-level AUC and AUPR of 0.9974 and 0.9986, respectively. Rockfish performed best among the other methods, with average AUC and AUPR of 0.9956 and 0.9978 (Fig. 5a). Unimeth also showed consistently high read-level performance across CpG density groups, genic and intergenic regions, and repetitive regions (Supplementary Fig. 18).

**Fig. 5.**
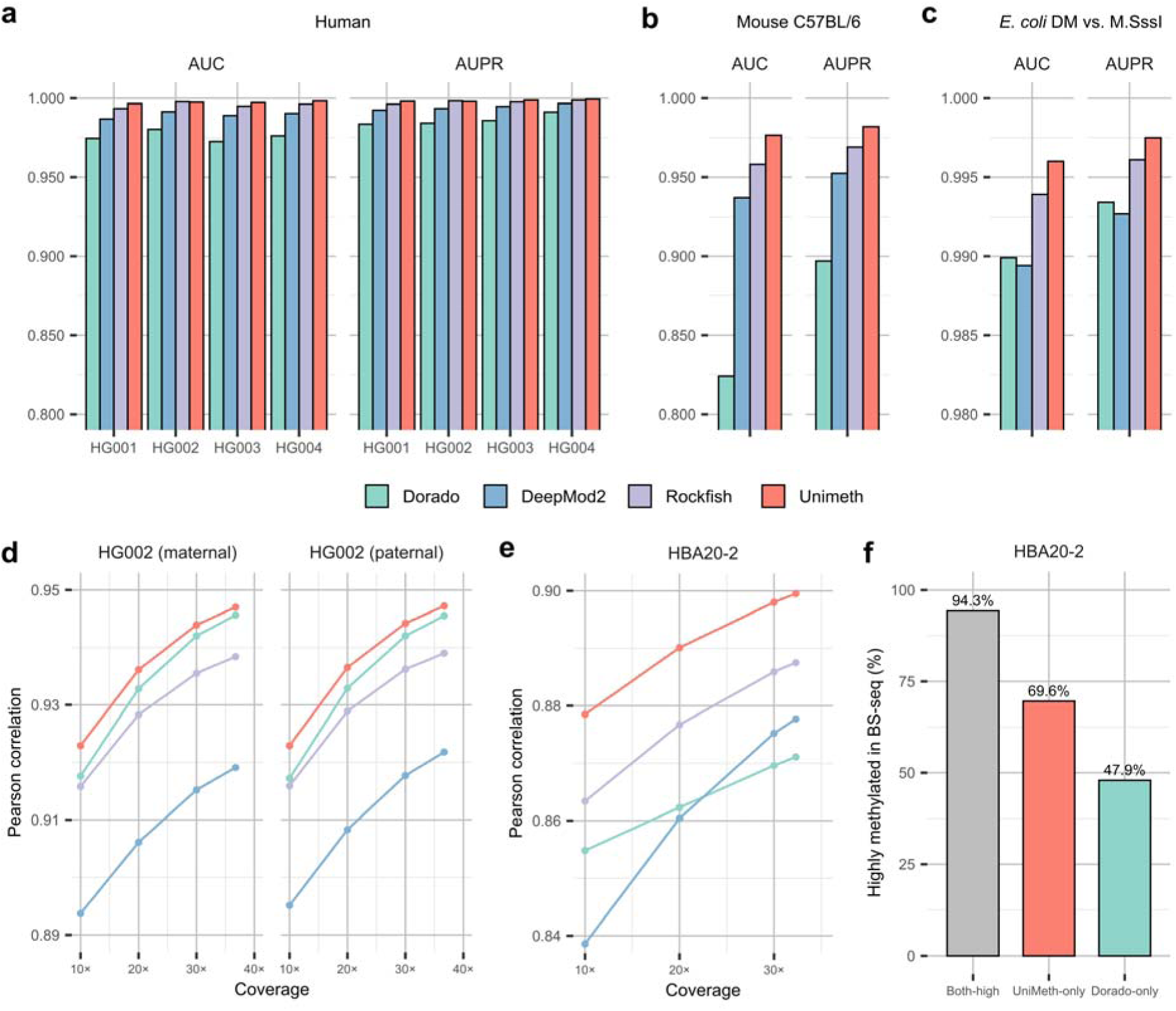
Evaluation of 5mCpG detection using nanopore R10.4.1 5kHz reads. **a** Read-level AUC and AUPR for Unimeth, Dorado, Rockfish and DeepMod2 on chromosome 1 from human samples HG001-HG004. **b,c** Read-level AUC and AUPR on mouse C57BL/6 (**b**) and *E. coli dam*/*dcm* methylase-knockout (DM) versus M.SssI-treated samples (**c**). **d** Site-level Pearson correlations with BS-seq for the maternal and paternal haplotypes of HG002 across nanopore read coverages. **e** Site-level Pearson correlations with BS-seq for the Alzheimer’s disease brain sample HBA20-2 across nanopore read coverages. In **d,e**, values at subsampled coverage levels represent the mean of five subsampling replicates. **f** Percentages of sites also classified as highly methylated by BS-seq among sites called highly methylated by both Unimeth and Dorado, by Unimeth only or by Dorado only.

We next evaluated the human-trained Unimeth model on mouse C57BL/6 and *E. coli* R10.4.1 5kHz datasets^25,27^. Unimeth achieved AUCs of 0.9766 in mouse and 0.9960 in the *E. coli dam*/*dcm* methylase-knockout (DM) versus M.SssI-treated comparison (Fig. 5b,c). These results were consistent across additional read-level metrics (Supplementary Fig. 19). Across the four human datasets and the mouse dataset, the average false-positive rates were 0.99% for Unimeth and 1.21% for Rockfish. The Unimeth false-positive rate was near zero in *E. coli* (Supplementary Fig. 22). We also trained Unimeth models for R10.4.1 4kHz and R9.4.1 reads. Both models showed high read-level performance on human datasets (Supplementary Figs. 20 and 21).

At the site level, we aggregated read-level CpG calls and compared methylation frequencies with matched enzymatic methyl-sequencing (EM-seq)^8^ or BS-seq data. Unimeth showed high correlations in human datasets from all three chemistries and mouse R10.4.1 5kHz data, including at lower coverages (Supplementary Figs. 23-25 and Supplementary Data 8-10). Using parental genotype information, we further assigned HG002 reads to maternal and paternal haplotypes (Methods). Unimeth achieved the highest Pearson correlations with BS-seq for both haplotypes across the read coverages (Fig. 5d and Supplementary Data 11). Using all available reads, Pearson correlations between Unimeth and BS-seq methylation frequencies were 0.9470 and 0.9472 for the maternal and paternal haplotypes, respectively.

To further validate Unimeth, we collected an Alzheimer’s disease brain sample, HBA20-2, and generated nanopore and matched BS-seq data (Methods, Supplementary Data 1-3). Unimeth achieved the highest Pearson correlations with BS-seq across all evaluated read coverages, reaching nearly 0.90 using all available reads (Fig. 5e and Supplementary Data 12). We next compared highly methylated CpG sites called by the four methods (Methods, Supplementary Fig. 26). Among sites called highly methylated by both Unimeth and Dorado, 94.3% were also highly methylated in BS-seq. The corresponding percentages were 69.6% for Unimeth-only sites and 47.9% for Dorado-only sites (Fig. 5f). The Unimeth-only groups also had higher percentages of sites highly methylated in BS-seq than the Rockfish-only and DeepMod2-only groups (Supplementary Fig. 27).

### Unimeth improves 6mA detection while retaining biological signals

6mA is the predominant DNA methylation mark in prokaryotes but occurs at much lower abundance in most eukaryotes^34,35^. By using a 6mA methyltransferase to label accessible DNA, Fiber-seq enables genome-wide profiling of chromatin accessibility and nucleosome positioning at single-molecule resolution^36,37^, while also providing a useful framework for benchmarking 6mA detection. We trained Unimeth on HG002 ONT Fiber-seq data^36^ and evaluated it on held-out chromosome 1 from the Fiber-seq and matched native genomic DNA (gDNA) datasets (Methods). This comparison assessed whether Unimeth could distinguish experimentally introduced 6mA signals from the lower background in native DNA. Across methylation-calling thresholds, Unimeth consistently reported lower fractions of methylated adenines than Dorado in both datasets (Fig. 6a). The estimates of Unimeth remained closer to orthogonal reference estimates, including 14.67% measured by tandem mass spectrometry (MS/MS) for Fiber-seq and 0.23% estimated by *fibertools*^36^ from PacBio data for native gDNA. For nucleosome profiling, we used *fibertools* to infer nucleosome lengths from Unimeth and Dorado ONT 6mA calls, with PacBio Fiber-seq nucleosome calls as a reference. Both methods produced peaks near the expected nucleosome size of 147 bp, with a more pronounced peak obtained for Unimeth (Fig. 6b).

**Fig. 6.**
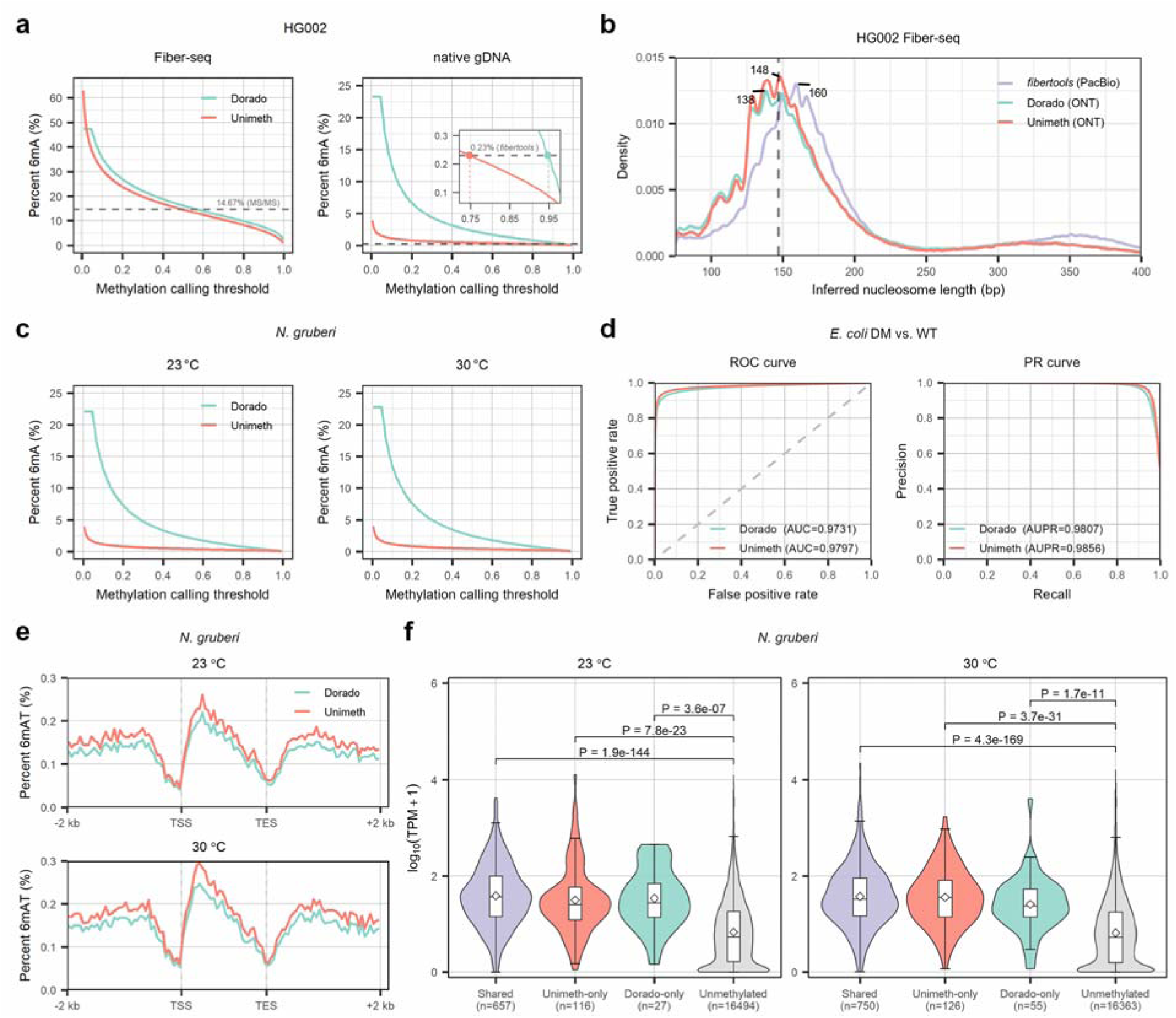
Evaluation of Unimeth for DNA 6mA detection using nanopore R10.4.1 5kHz reads. **a** Percent 6mA as a function of methylation calling threshold in the nanopore data of HG002 Fiber-seq and native gDNA; dashed reference lines indicate orthogonal estimates (14.67% from MS/MS for Fiber-seq and 0.23% from *fibertools* for native gDNA); reads aligned to chromosome 1 were used for testing. **b** Density of inferred nucleosome lengths in the HG002 Fiber-seq datasets. Nucleosome lengths were inferred by *fibertools* from ONT 6mA calls generated by Unimeth and Dorado, and compared with *fibertools* calls from PacBio Sequel II data; 10,000 reads were sampled from chromosome 1 for testing. **c** Percent 6mA as a function of methylation calling threshold in the *N. gruberi* samples grown at 23 °C and 30 °C. **d** Receiver operating characteristic (ROC) and precision–recall (PR) curves for read-level 6mA detection on the *E. coli* dataset at GATC sites. **e** Metaplots of 6mAT levels across scaled gene regions in *N. gruberi* samples. **f** Gene expression distributions in *N. gruberi* samples. Boxes show interquartile ranges with median lines; whiskers extend to the most extreme values within 1.5× the interquartile range from the box edges, and white diamonds indicate means. P values were calculated using two-sided Wilcoxon rank-sum tests.

We evaluated read-level 6mA detection at GATC sites using an independent *E. coli* nanopore dataset from wild-type (WT) and *dam/dcm* methylase-knockout (DM) strains^27^. Across 3.9 million read-level GATC sites, Unimeth achieved an AUC of 0.9797 and an AUPR of 0.9856, compared with 0.9731 and 0.9807 for Dorado, respectively (Fig. 6d).

We also evaluated Unimeth on nanopore data from *N. gruberi* cultures grown at 23 °C and 30 °C^38^. Unimeth consistently called fewer 6mA sites than Dorado across methylation-calling thresholds at both temperatures, suggesting lower background calling (Fig. 6c). Following Charria *et al.*^38^, we applied a probability threshold of 0.995 to define methylated 6mA sites for downstream analysis (Methods). At ApT sites (6mAT), Unimeth preserved clear metagene-scale patterns, including depletion around transcription start sites (TSSs) and transcription end sites (TESs). Unimeth 6mAT profiles also showed enrichment immediately downstream of TSSs, consistent with reported 6mA enrichment downstream of TSSs in expressed genes^38^ (Fig. 6e). Grouping genes by expression level further showed that 6mAT signals became progressively more pronounced in more highly expressed genes at both temperatures (Supplementary Fig. 28). At the gene level, Unimeth identified more 6mA-methylated genes than Dorado at both temperatures (Supplementary Fig. 29). Genes called methylated by both methods or by Unimeth alone showed significantly higher expression than unmethylated genes at both temperatures (Fig. 6f). Together, these analyses indicate that Unimeth reduces background 6mA calls while retaining signals for nucleosome profiling and gene-level analysis across the evaluated datasets.

## Discussion

We developed Unimeth as a unified framework for 5mC and 6mA detection from nanopore reads across species and sequencing chemistries. Unimeth combines patch-based joint methylation calling with a multi-phase training procedure. Ablation analyses showed that the training phases had different effects. Pre-training improved read-level discrimination, whereas calibration improved site-level methylation quantification, particularly for CHH sites.

We evaluated Unimeth using public and in-house datasets covering diverse species and sample types. In plants, Unimeth improved 5mC detection in non-CpG contexts, where methylated cytosines are often less abundant and false-positive calls can substantially affect methylation estimates. Reduced background calls were consistently observed at high-confidence unmethylated sites and in methylation-deficient *A. thaliana* mutants. Unimeth also achieved high 5mCpG performance in human and mouse datasets. Validation in an Alzheimer’s disease brain sample further supported accurate CpG methylation profiling with Unimeth. For 6mA, Unimeth reduced background calls in native gDNA and accurately distinguished *E. coli* WT and DM reads at GATC sites. Its 6mA profiles also supported nucleosome analysis in Fiber-seq data and showed gene-level associations with expression in *N. gruberi*.

The reference datasets assessed different aspects of methylation detection. Matched BS-seq or EM-seq data enabled quantitative assessment of site-level methylation frequencies. Methylation-deficient mutants and native gDNA provided low-methylation controls for 5mC and 6mA, respectively. Enzyme-treated bacterial samples and Fiber-seq data provided methylation-enriched comparisons.

Unimeth is an open and reusable framework for nanopore methylation calling. The training and testing sets are explicitly defined, and the corresponding data, code, pretrained models and training workflow are publicly available. Together with standard nanopore input and output formats, these resources should facilitate reproducible benchmarking and future adaptation to additional datasets, chemistries or DNA modifications.

Unimeth currently relies on basecalled sequences and signal-to-sequence mappings produced by existing basecalling workflows. Because basecallers such as Dorado are actively updated, model performance may vary with basecalling models and sequencing chemistries. Future methods could reduce this dependence by learning more transferable representations directly from raw nanopore signals.

## Methods

### Preparation of plant materials and DNA extraction

*Arabidopsis thaliana (L.) Heynh. Columbia-0* (Col-0), the *A. thaliana* mutants *ddcc*^31^ and *met1-3* (CS16394)^39^, *Oryza sativa L. ssp. japonica cv. Nipponbare*, and *Musa* spp. were used in this study. All *A. thaliana* lines were in the Col-0 background. *A. thaliana* seeds were surface-sterilized in 4% sodium hypochlorite, stratified for 2 days at 4 °C, and grown under long-day conditions (16 h light/8 h dark, 22 °C). Leaves were collected from 21-day-old Col-0 wild-type plants, whereas aerial tissues were collected from *ddcc* and *met1-3* mutant seedlings. *O. sativa* seeds were sown in soil and grown for 28 days in a growth chamber at 28 °C and 70% relative humidity under a 16 h/8 h light/dark cycle. *O. sativa* leaves were harvested, immediately frozen in liquid nitrogen, and stored at −80 °C until use. *Musa* spp. material was provided by Wuhan Frasergen Bioinformatics Co., Ltd. Genomic DNA was extracted using a QIAGEN genomic DNA extraction kit according to the manufacturer’s instructions. DNA purity was assessed using a NanoDrop One UV-Vis spectrophotometer (Thermo Fisher Scientific).

### Ethics statement and human sample collection

This study was conducted in accordance with applicable Chinese regulations on human genetic resources. The collection and sequencing of HBA20-2 were approved by the Ethics Committee of Xiangya Hospital, Central South University (No. 2021111238) and conducted in accordance with the Declaration of Helsinki. Brain tissue sample HBA20-2 was collected from the dorsolateral prefrontal cortex (DLPFC) of a donor with Alzheimer’s disease. Written informed consent was obtained from the donor’s legally authorized representatives. Genomic DNA for nanopore sequencing and BS-seq was extracted by proteinase K digestion followed by phenol-chloroform-isoamyl alcohol extraction and isopropanol precipitation.

### Nanopore sequencing and BS-seq

We generated nanopore data for *A. thaliana* Col-0 and the *ddcc* and *met1-3* mutants, *O. sativa* cv. Nipponbare, *Musa* spp. and the human brain sample HBA20-2. Matched BS-seq data were also generated for all samples except the two *A. thaliana* mutants. For nanopore sequencing, genomic DNA was size-selected using the BluePippin system (Sage Science). Sequencing adapters were ligated using SQK-LSK114 for plant samples and SQK-LSK114-XL for HBA20-2 (ONT). Libraries were quantified using a Qubit 4.0 Fluorometer (Invitrogen) and sequenced on a PromethION instrument using R10.4.1 flow cells (FLO-PRO114M) at 5 kHz.

For BS-seq, genomic DNA was subjected to bisulfite conversion, followed by library preparation. For plant samples, bisulfite conversion and library preparation were performed using the TIANGEN DNA Bisulfite Conversion Kit (Cat#DP215, TIANGEN BIOTECH) and TruSeq DNA Methylation Kit (Cat#EGMK91324, Illumina), respectively. For HBA20-2, the corresponding kits were the EZ DNA Methylation-Gold Kit (Zymo Research) and Accel-NGS Methyl-Seq DNA Library Kit (Swift, Cat#30096). After PCR amplification, libraries were sequenced on a NovaSeq 6000 instrument (Illumina) to generate paired-end 150-bp reads.

### Public nanopore, BS-seq, EM-seq and PacBio sequencing data

We collected publicly available nanopore sequencing datasets of 13 organisms, including 9 plants (*A. thaliana*, *B. vulgaris*, *C. sinensis*, *O. sativa*, *R. communis*, *S. lycopersicum*, *S. miltiorrhiza*, *S. tuberosum*, and *V. vinifera*), 2 mammals (*H. sapiens* and *M. musculus*), 1 heterolobosean protist (*N. gruberi*), and 1 bacterium (*E. coli*), generated using ONT R10.4.1 5kHz, R10.4.1 4kHz and R9.4.1 chemistries.

For the R10.4.1 5kHz chemistry, we used nanopore reads from the 9 plant species reported by Chen *et al.*^26^, 4 *H. sapiens* samples (HG001-HG004) from the ONT Open Data project (https://epi2me.nanoporetech.com/dataindex/), HG002 Fiber-seq and matched native gDNA data from Jha *et al*.^36^, *M. musculus* C57BL/6 from Stanojević *et al.*^25^, and *N. gruberi* cultured at 23 °C and 30 °C from Charria *et al.*^38^. We also used *E. coli* nanopore data from Kulkarni *et al.*^27^, including reads from a wild-type strain (WT), a *dam/dcm* methylase-knockout strain (DM), and an M.SssI-treated sample (M.SssI). For the R10.4.1 4kHz chemistry, we used nanopore reads of 3 *H. sapiens* samples (HG002-HG004) from the ONT Open Data project. For the R9.4.1 chemistry, we used nanopore reads of HG002-HG004 from the ONT Open Data project, HG001 from Jain *et al.*^40^ (see Supplementary Data 13), and *A. thaliana* and *O. sativa* from our previous study^29^.

To benchmark Unimeth for 5mC detection, we also collected the corresponding BS-seq or EM-seq datasets. For the 9 plant species, the matched BS-seq data were obtained from Chen *et al.*^26^ and Ni *et al.*^29^. For *H. sapiens*, we used BS-seq data for HG002 from the ONT Open Data project and EM-seq data for HG001, HG003 and HG004 from the SEQC2 study^11^. For *M. musculus*, the matched BS-seq data were obtained from Stanojević *et al.*^25^. In addition, we used PacBio Sequel II Fiber-seq data of HG002 from Jha *et al*.^36^ as an orthogonal reference for evaluating nucleosome profiles inferred from ONT 6mA calls.

### Reference genomes and annotations

We used Col-CEN^13^ and T2T-NIP^41^ (AGIS1.0) as the reference genomes for *A. thaliana* and *O. sativa*, respectively, together with the corresponding gene and centromere annotations. For *Musa* spp., we generated a reference assembly using hifiasm^42^ (v0.25.0-r726) and purge_dups^43^ (v1.2.6). For the other 7 plant species, we used the same reference genomes as Chen *et al.*^26^. For the publicly available human datasets, we used CHM13 v2.0^44^, with gene annotation from the GitHub repository marbl/CHM13 (https://github.com/marbl/CHM13). CpG island, peri/centromeric satellite and RepeatMasker annotations for CHM13 were obtained from UCSC (T2T CHM13v2.0/hs1). For HBA20-2, we used the GRCh38 primary assembly from GENCODE^45^. For *M. musculus*, we used the GRCm39 (Release M37) genome from GENCODE^45^. For *E. coli*, we used the NCBI reference genome GCF_000005845.2. For *N. gruberi*, we used the re-assembled genome and corresponding gene annotation reported by Charria *et al.*^38^. Details of all reference genomes and annotations are provided in Supplementary Note 1.

### Processing of BS-seq and EM-seq data

All BS-seq and EM-seq datasets were processed using the nf-core/methylseq pipeline^46^ (v2.5.0). We ran the Bismark^47^ workflow in methylseq with the parameters “--comprehensive true --cytosine_report true” to obtain strand-specific methylation information for all three sequence contexts (CpG, CHG, and CHH). For HBA20-2, we used “--ignore_r2 12 --ignore_3prime 2 --ignore_3prime_r2 2” during Bismark methylation extraction to reduce read-end methylation bias. Per-site methylation frequencies were calculated from Bismark methylation calls and saved in bedmethyl format. For samples with multiple replicates (*H. sapiens* HG001, HG003 and HG004; *M. musculus* C57BL/6; and *A. thaliana* Col-0; see Supplementary Data 3), we used the average methylation frequencies across replicates as the final results. Only sites covered by at least five reads in all replicates were retained for further evaluation. Details of the BS-seq and EM-seq datasets used in this study, including read depth and methylation levels, are provided in Supplementary Data 3.

### Processing of nanopore data

The nanopore datasets used in this study included R10.4.1 5kHz, R10.4.1 4kHz and R9.4.1 reads in POD5 or FAST5 format. For 5mC detection, we used Dorado (v0.7.1) to basecall all the nanopore raw reads. To retain signal-to-sequence alignment information, we enabled the “--emit-moves” option during basecalling. Raw reads in FAST5 format were converted to POD5 format before basecalling. We used the models “dna_r10.4.1_e8.2_400bps_sup@v5.0.0”, “dna_r10.4.1_e8.2_400bps_sup@v4.1.0”, and “dna_r9.4.1_e8_sup@v3.3” for R10.4.1 5kHz, R10.4.1 4kHz and R9.4.1 reads, respectively. For HBA20-2, only reads with a mean quality score ≥10 were retained for downstream analyses. For 6mA detection in the *E. coli* WT and DM datasets, HG002 Fiber-seq and matched native gDNA datasets, and *N. gruberi* datasets, we used Dorado (v0.9.1) with the model “dna_r10.4.1_e8.2_400bps_sup@v5.0.0_6mA@v3” for basecalling. Dorado integrates minimap2^48^ to perform read-to-reference alignment during basecalling, and the outputs were saved in BAM format. Details of the nanopore datasets used in this study, including sequencing chemistry, number of reads, read length, read depth and read quality, are provided in Supplementary Data 1. Reads used for model training and evaluation were split by chromosome or contig (Supplementary Data 2).

To benchmark Unimeth for 5mC detection, we compared it with Dorado (v0.7.1), Rockfish (v0.3), DeepMod2 (v0.3.1), DeepPlant (v1.1.0) and DeepSignal-plant (v0.1.6). Dorado methylation calling was performed during basecalling. For R10.4.1 5kHz, R10.4.1 4kHz and R9.4.1 reads, we used the Dorado modified-base models “dna_r10.4.1_e8.2_400bps_sup@v5.0.0_5mC_5hmC@v1”, “dna_r10.4.1_e8.2_400bps_sup@v4.1.0_5mCG_5hmCG@v2”, and “dna_r9.4.1_e8_sup@v3.3_5mCG@v0.1”, respectively. We ran Rockfish with the models “rf-r10.4.1-5kHz-base.pt” and “rf-base.pt” to call CpG methylation from R10.4.1 5kHz and R9.4.1 reads, respectively. Before calling methylation from R9.4.1 reads with Rockfish, reads were basecalled using Guppy (v6.2.1) with the model configuration “dna_r9.4.1_450bps_sup_prom.cfg”. For DeepMod2, we used the models “bilstm_r10.4.1_5khz_v5.0”, “bilstm_r10.4.1_4khz_v4.1”, and “bilstm_r9.4.1” to call CpG methylation from R10.4.1 5kHz, R10.4.1 4kHz and R9.4.1 reads, respectively. We used DeepPlant with the default R10.4.1 5kHz model and parameters to call 5mC methylation in plant nanopore datasets. Following Chen *et al.*^26^, reads were first basecalled using Dorado with the model “dna_r10.4.1_e8.2_400bps_hac@v5.0.0” before DeepPlant analysis. We used DeepSignal-plant (v0.1.6) with the recommended pipeline, including Guppy (v6.4.2) for basecalling and Tombo (v1.5.1) for re-squiggle, to call 5mC methylation from R9.4.1 reads of *A. thaliana* and *O. sativa*.

### Unimeth model architecture

Unimeth takes raw nanopore signals and their corresponding basecalled sequences as input. For R10.4.1 reads, we obtained the signal-to-sequence alignments based on the move tables from the BAM files. For R9.4.1 reads, we re-squiggled the signals to reference genomes (Supplementary Note 2). During pre-training, sequences consisted of nucleotide tokens only. During fine-tuning, calibration and inference, special context tokens ([CpG], [CHG] and [CHH] for 5mC detection, and [A] for 6mA detection) were inserted after the corresponding target bases. Sequences were divided into overlapping 256-token sequence segments, and each segment was paired with the corresponding signal segment using the signal-to-sequence mapping table. We refer to each paired signal-sequence segment as a patch. During inference, a single prediction was retained for each target site to avoid duplicate calls from overlapping patches.

The model architecture consists of an encoder and a decoder. The encoder is designed to learn representations from raw signal inputs and comprises three core components: a signal processor, a positional encoder and a transformer backbone. The signal processor normalizes the raw signals and uses two convolutional neural network (CNN) layers to convert them into signal embeddings. The positional encoder adds positional information to the signal embeddings using sinusoidal positional encoding, which can be formulated as:

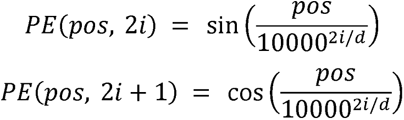

where *pos* denotes the signal position, *i* is the dimension index, and *d* is the embedding dimension. The signal and positional embeddings are combined and passed through 12 transformer encoder layers to produce the final signal embeddings.

The decoder processes the tokenized DNA sequence with inserted context tokens. It consists of a token embedding layer, positional embeddings and a transformer backbone. The tokenizer converts the sequence and special tokens into token embeddings, to which positional embeddings are added. The decoder backbone contains 12 transformer decoder layers that attend to both the token embeddings and the signal embeddings from the encoder.

During pre-training, the decoder reconstructs the nucleotide sequence within each patch. During fine-tuning and methylation calling, it maintains the patch structure and replaces each methylation token with a prediction token, either “[+]” indicating a methylated site or “[-]” indicating an unmethylated site.

### Model training

Unimeth was trained through up to three phases: pre-training, fine-tuning, and calibration.

#### Pre-training

During pre-training, the model learned signal-sequence representations by reconstructing the nucleotide sequences corresponding to raw nanopore signals, without inserting methylation context tokens. Training used a next-token prediction objective with standard cross-entropy loss:

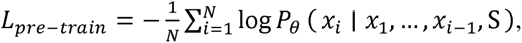

where *x_i_* is the nucleotide at position *i*, *S* is the corresponding signal patch, *N* is the sequence length and θ denotes the model parameters. The resulting weights were used to initialize subsequent methylation fine-tuning.

#### Fine-tuning

During fine-tuning, context tokens were inserted into the decoder sequence to mark the methylation context and target position. For example, the sequence “C G” at a CpG site was transformed into “C [CpG] G”. The causal mask used during pre-training was removed to allow the decoder to use the complete sequence context. At each marked position, the model predicted either “[+]” (methylated) or “[−]” (unmethylated), producing outputs such as “C [+] G” or “C [−] G”.

For 5mC models trained using quantitative site-level reference measurements (BS-seq or EM-seq), target cytosines in nanopore reads mapped to sites with methylation frequencies below 20% were assigned unmethylated labels, whereas those mapped to sites with frequencies above 80% were assigned methylated labels. Target cytosines mapped to sites with intermediate methylation frequencies were excluded from the initial fine-tuning phase. For models using a frequency-weighted loss, the weight assigned to a labelled position was:

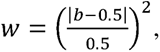

where *b* is the reference methylation frequency. The loss was calculated only at positions marked by methylation context tokens. For plant 5mC models, cross-entropy losses were calculated separately for methylated and unmethylated CpG, CHG and CHH observations and then summed. For models with a single target context, weighted cross-entropy was used. For 6mA fine-tuning, binary pseudo-labels for individual adenines were derived from Dorado probabilities rather than site-level methylation-frequency thresholds, as described below.

#### Calibration

When quantitative site-level references were available, calibration was used to incorporate sites across the full range of methylation frequencies. Genomic sites covered by more than five nanopore reads were retained for calibration. For each site *s*, the target number of methylated reads 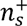 was calculated from the reference methylation frequency ρ*_s_* and the total number of reads covering that site *N_s_*:

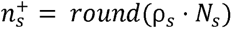

The fine-tuned model was used to predict a methylation probability *p_s,i_* ∊ [0,1] for read *i* at site *s*. Reads covering each site were then ranked in descending order of their predicted probabilities:

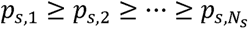

The top 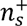 observations were assigned methylated pseudo-labels, and all other observations were assigned unmethylated pseudo-labels:

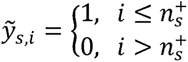

This assignment made the proportion of methylated pseudo-labels approximate the reference methylation frequency as closely as permitted by nanopore coverage. The pseudo-labels were used for additional fine-tuning on reads from the training chromosomes or contigs.

#### Training datasets and parameters

During pre-training, separate models were trained for the R9.4.1, R10.4.1 4kHz and R10.4.1 5kHz chemistries using the corresponding human nanopore datasets (Supplementary Data 2). Task-specific models were subsequently developed for different modification types and sequencing chemistries.

For plant 5mC detection, the R10.4.1 5kHz model was trained on *R. communis*, *S. miltiorrhiza* and *S. tuberosum*, following the training dataset used in DeepPlant^26^. The R9.4.1 model was trained on *A. thaliana* and *O. sativa*, following the experimental setup of DeepSignal-plant^29^. For mammalian 5mCpG detection, three chemistry-specific models were fine-tuned on HG002 R9.4.1, R10.4.1 4kHz and R10.4.1 5kHz data, respectively. To assess the effect of patch length, we fine-tuned models from random initialization using patch lengths of 64, 128 and 256 tokens on R10.4.1 5kHz data from *R. communis*, *S. miltiorrhiza* and *S. tuberosum*. The results showed that read-level performance generally improved with increasing patch length (Supplementary Fig. 30).

For 6mA detection, a chemistry-specific model was trained using HG002 ONT Fiber-seq data generated with R10.4.1 5kHz chemistry. Chromosome 1 was excluded from training and used for testing. Dorado probabilities were used to define binary pseudo-labels for individual adenine observations. Adenine observations with 6mA probabilities of at least 0.95 were treated as positive.

Observations with probabilities of 0 within predicted nucleosome regions were treated as negative. Two additional models were trained using positive pseudo-label thresholds of 0.90 and 0.99, with all other settings unchanged. The three models were evaluated at read-level GATC sites using the independent *E. coli* WT/DM benchmark (Supplementary Fig. 31). Only pre-training and fine-tuning were performed for the 6mA model.

The training and testing data for each model are listed in Supplementary Data 2. Detailed training parameters are provided in Supplementary Note 3.

### Evaluation on 5mC detection

#### Evaluation at read level

We used the final calibrated Unimeth models for comparisons with other methods at both the read and site levels. For *H. sapiens*, *M. musculus* and plant samples, high-confidence unmethylated cytosine sites were defined from BS-seq or EM-seq data as sites with methylation frequency = 0 and coverage ≥ 5. For samples with multiple replicates, all replicates were required to meet these criteria. High-confidence methylated cytosine sites were defined as sites with methylation frequency = 100% and coverage ≥ 5 for samples with a single replicate, or methylation frequency ≥ 95% and coverage ≥ 5 in all replicates for multi-replicate samples. Positive and negative read-level samples were then extracted from nanopore reads covering these sites. Unless otherwise specified, Unimeth probabilities ≥ 0.5 were classified as methylated and probabilities < 0.5 as unmethylated. Accuracy, sensitivity, specificity, precision and FPR were calculated from binary calls, whereas AUC and AUPR were calculated from methylation probabilities. For the *E. coli* dataset, positive and negative samples were extracted from M.SssI-treated and *dam/dcm* methylase-knockout reads, respectively. For methylation-deficient mutant analyses, methylation levels were calculated in 10-kb windows separately for CpG, CHG and CHH contexts as methylated calls divided by total covered calls within each window.

#### Evaluation at site level

We evaluated Unimeth at the genome-wide site level by comparing per-site methylation frequencies predicted by Unimeth with BS-seq or EM-seq results. Pearson correlation (*r*), coefficient of determination (*r*^2^), Spearman correlation (ρ), and root mean square error (RMSE) were calculated from per-site methylation frequencies. For each comparison, only sites with at least 5× coverage in both nanopore and BS-seq or EM-seq data were retained. To assess performance under different nanopore coverages, nanopore reads were randomly subsampled using rasusa^49^ (v2.0.0).

For HG002, we used parental phasing results from the Illumina WGS trio analysis reported by our previous study^15^. We used WhatsHap^50^ (v2.8) to assign nanopore reads to maternal and paternal haplotypes. Haplotype-specific CpG methylation frequencies were compared with the corresponding BS-seq results.

For HBA20-2, we compared highly methylated CpG sites (methylation frequency ≥80% and coverage ≥10) in BS-seq and for each of the four nanopore methods. Unimeth was compared separately with Dorado, Rockfish and DeepMod2, grouping sites as highly methylated by both methods or by either method alone. For each group, we calculated the percentage of sites also highly methylated in BS-seq.

### Evaluation and downstream analysis of 6mA detection

We evaluated Unimeth for 6mA detection using R10.4.1 5kHz nanopore datasets from HG002 ONT Fiber-seq and matched native gDNA, *E. coli* wild-type (WT) and *dam/dcm* methylase-knockout mutant (DM) samples, and *N. gruberi* samples cultured at 23 °C and 30 °C. For HG002, chromosome 1 Fiber-seq and matched native gDNA data were used to assess 6mA predictions across methylation-calling thresholds, and nucleosome lengths inferred from ONT 6mA calls were compared with nucleosome profiles from PacBio Fiber-seq data. For *E. coli*, read-level predictions at GATC sites were evaluated using AUC and AUPR. For *N. gruberi*, we followed Charria *et al.*^38^ and used a probability threshold of 0.995 to define methylated 6mA sites for downstream analyses focused on ApT sites, including metagene-scale profiling, expression-based gene grouping, and comparison of gene expression between methylation classes using the Wilcoxon rank-sum test. Detailed methods and parameters are provided in Supplementary Note 4.

## Data availability

The ONT R10.4.1 5kHz and BS-seq reads generated in this study for *A. thaliana* Col-0, *O. sativa* and *Musa* spp., together with ONT reads from the *ddcc* and *met1-3* mutants, have been deposited in NGDC (National Genomics Data Center) under BioProject accession PRJCA067807 [https://ngdc.cncb.ac.cn/bioproject/browse/PRJCA067807] and GSA (Genome Sequence Archive) accession CRA045731 [https://ngdc.cncb.ac.cn/gsa/browse/CRA045731]. The ONT R10.4.1 5kHz and matched BS-seq data for HBA20-2 will be deposited in GSA-Human at NGDC. The data will be available under controlled access through GSA-Human, subject to approval by the Data Access Committee (DAC).

The R10.4.1 5kHz nanopore reads of *A. thaliana*, *B. vulgaris*, *C. sinensis*, *O. sativa*, *R. communis*, *S. lycopersicum*, *S. miltiorrhiza*, *S. tuberosum*, and *V. vinifera* generated by Chen *et al*.^26^, together with the corresponding BS-seq reads, are available from the BioProjects of NGDC under accessions PRJCA030666 [https://ngdc.cncb.ac.cn/bioproject/browse/PRJCA030666] and PRJCA023349 [https://ngdc.cncb.ac.cn/bioproject/browse/PRJCA023349]. The R9.4.1 nanopore reads of *A. thaliana* and *O. sativa*, together with the corresponding BS-seq reads, are available from the BioProject of NGDC under accession PRJCA004326 [https://ngdc.cncb.ac.cn/bioproject/browse/PRJCA004326]. The R9.4.1 nanopore reads of *A. thaliana* Col-0, *ddcc* and *met1-3* are available from the BioProject of NGDC under accession PRJCA013292 [https://ngdc.cncb.ac.cn/bioproject/browse/PRJCA013292].

The R10.4.1 5kHz nanopore reads of HG001, HG002, HG003, and HG004 are available from the ONT Open Data project [https://epi2me.nanoporetech.com/giab-2023.05/]. The R10.4.1 4kHz nanopore reads of HG002, HG003, and HG004 are available from the ONT Open Data project [https://epi2me.nanoporetech.com/askenazi-kit14-2022-12/]. The R9.4.1 nanopore reads of HG001 are available from the GitHub repository [https://github.com/nanopore-wgs-consortium/NA12878], and the R9.4.1 nanopore reads of HG002 are available from the ONT Open Data project [https://labs.epi2me.io/gm24385_2020.11/] with the flowcell ID PAG07165. The R9.4.1 nanopore reads of HG003 and HG004 are available from Human Pangenome Reference Consortium^51^ [https://s3-us-west-2.amazonaws.com/human-pangenomics/index.html?prefix=NHGRI_UCSC_panel/HG003/nanopore/ and https://s3-us-west-2.amazonaws.com/human-pangenomics/index.html?prefix=NHGRI_UCSC_panel/HG004/nanopore/]. The BS-seq reads of HG002 are available from the ONT Open Data project [https://labs.epi2me.io/gm24385-5mc/], and the EM-seq data of HG001, HG003, and HG004 are available under NCBI BioProject PRJNA200694 [https://www.ncbi.nlm.nih.gov/bioproject/PRJNA200694]. The HG002 ONT Fiber-seq and matched native gDNA datasets are available under NCBI BioProject PRJNA956114 [https://www.ncbi.nlm.nih.gov/bioproject/?term=PRJNA956114]. PacBio Sequel II Fiber-seq data of HG002 were provided by the authors of Jha *et al.*^36^ upon request.

The R10.4.1 5kHz nanopore reads of *M. musculus* C57BL/6 and the corresponding BS-seq reads from Stanojević *et al.*^25^ are available under NCBI BioProject PRJNA876781 [https://www.ncbi.nlm.nih.gov/bioproject/PRJNA876781/]. The R10.4.1 5kHz nanopore reads of *E. coli* from Kulkarni *et al*.^27^ are available from AWS [https://registry.opendata.aws/ont_basemod_data/]. The *N. gruberi* R10.4.1 5kHz nanopore reads and the RNA-seq datasets from Charria *et al.*^38^ are available under NCBI BioProject PRJNA1089232 [https://www.ncbi.nlm.nih.gov/bioproject/PRJNA1089232].

## Code availability

Unimeth is publicly available at GitHub [https://github.com/sekeyWang/Unimeth].

## Acknowledgements

This work was supported in part by the National Natural Science Foundation of China (grant Nos. 62332020, 62302526) and the Project of Xiangjiang Laboratory (No. 23XJ01011). We also acknowledge support from the Center for Computational Science and Engineering at Southern University of Science and Technology. We thank Onkar Kulkarni and Divya Tej Sowpati from CSIR Centre for Cellular and Molecular Biology for providing the *E. coli* ONT data. We thank Andrew B. Stergachis and Mitchell R. Vollger from University of Washington for providing the PacBio and ONT Fiber-seq data of HG002. We thank Jonathan Cahn and Evan Ernst from Cold Spring Harbor Laboratory for providing their ONT data of *A. thaliana* mutants and useful discussions. We thank Xiaowen Feng of Oxford Nanopore Technologies for her assistance. We also thank Pierre Baduel from Université Paris Sciences et Lettres, Kateryna Makova from Penn State University, and Brandon D. Pickett from National Human Genome Research Institute (NHGRI) for providing their data and useful discussions. This work was carried out in part using computing resources at the High-Performance Computing Center of Central South University.

## Author contributions

S.K.W., P.N., and J.X.W. conceived and designed this project. S.K.W., P.N., Y.F.X., and T.S. developed and evaluated Unimeth using the sequencing data. N.H. helped design the Unimeth model.

J.X.Z. and Y.S. generated the ONT data for the *A. thaliana* mutants and helped analyze the data.

H.D.L., L.S., and X.X.Y. collected and helped analyze the HBA20-2 sequencing data. F.L. helped design the model and the experiments. J.X.W. supervised the study. All authors have read and approved the final version of this paper.

