## Supplementary Information for "Unimeth: a unified transformer framework for accurate DNA methylation detection from nanopore reads"

Wang *et al.*

### Supplementary Figures

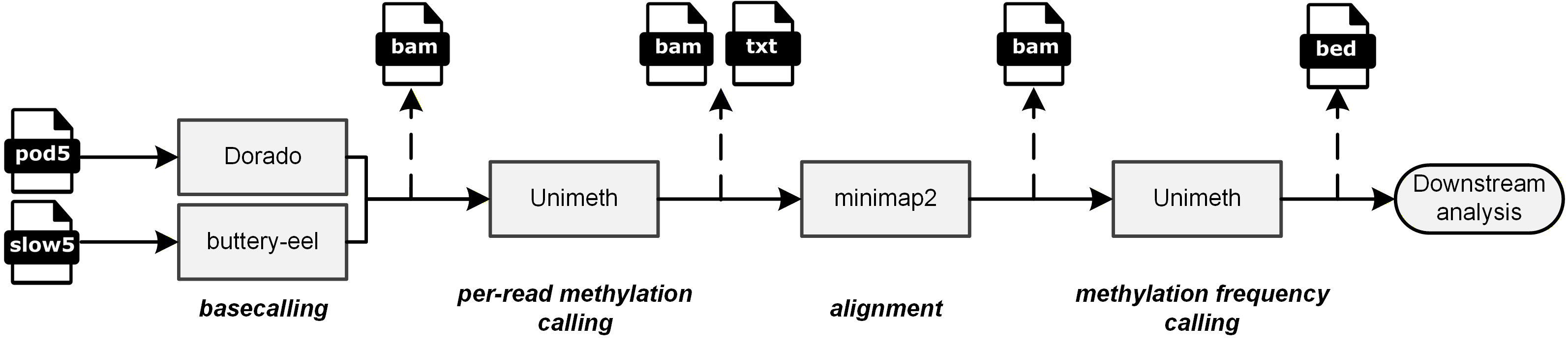

**Supplementary Fig. 1** Workflow of Unimeth for per-read and site-level methylation calling. POD5 or SLOW5 signal files and basecalled reads are used for per-read methylation calling. After alignment to a reference genome, Unimeth aggregates read-level calls to estimate site-level methylation frequencies for downstream analysis.

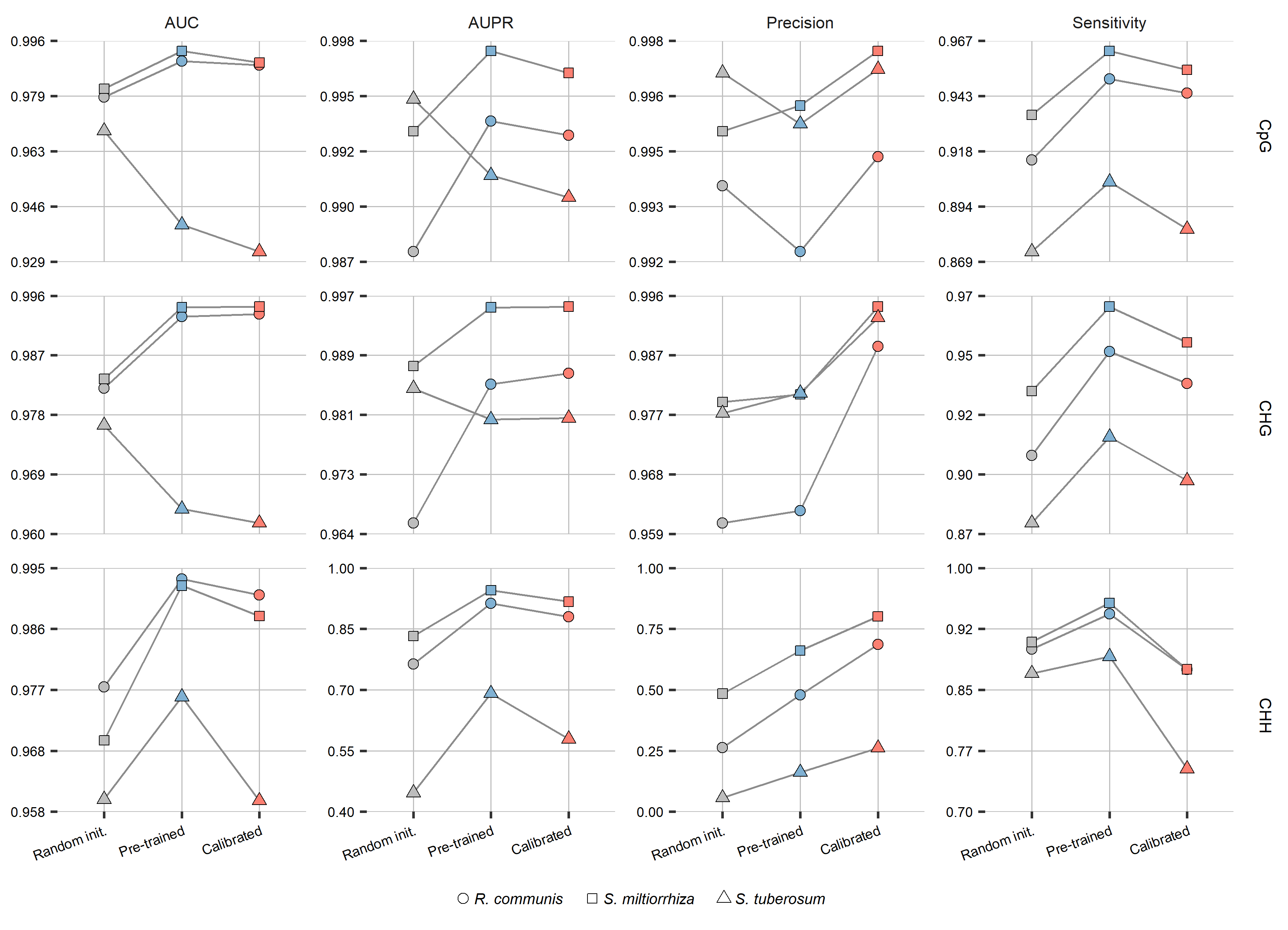

**Supplementary Fig. 2** Read-level effects of pre-training and calibration across cytosine methylation contexts. AUC, AUPR, precision and sensitivity are shown for CpG, CHG and CHH methylation in *R. communis*, *S. miltiorrhiza* and *S. tuberosum*, evaluated on held-out chromosomes or contigs. All model variants used the same fine-tuning data and procedure. Random init. started from random weights, Pre-trained started from pre-trained weights, and Calibrated included the calibration step.

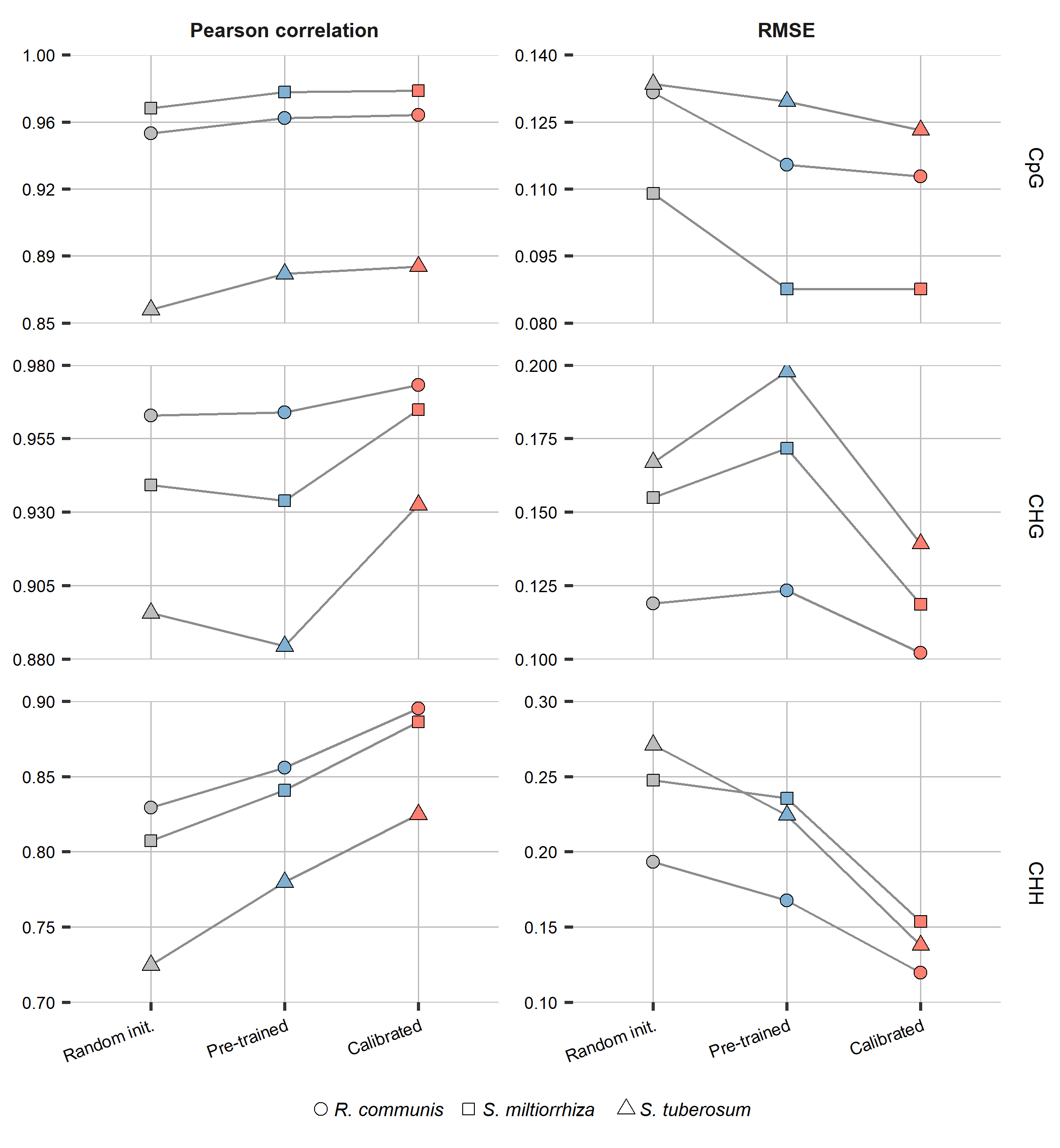

**Supplementary Fig. 3** Site-level effects of pre-training and calibration across cytosine methylation contexts. Pearson correlations and root mean square errors (RMSEs) between site-level methylation frequencies predicted by the three model variants and measured by BS-seq at 20× nanopore coverage are shown for CpG, CHG and CHH methylation in *R. communis*, *S. miltiorrhiza* and *S. tuberosum*. Values represent the mean of five subsampling replicates. All model variants used the same fine-tuning data and procedure. Random init. started from random weights, Pre-trained started from pre-trained weights, and Calibrated included the calibration step.

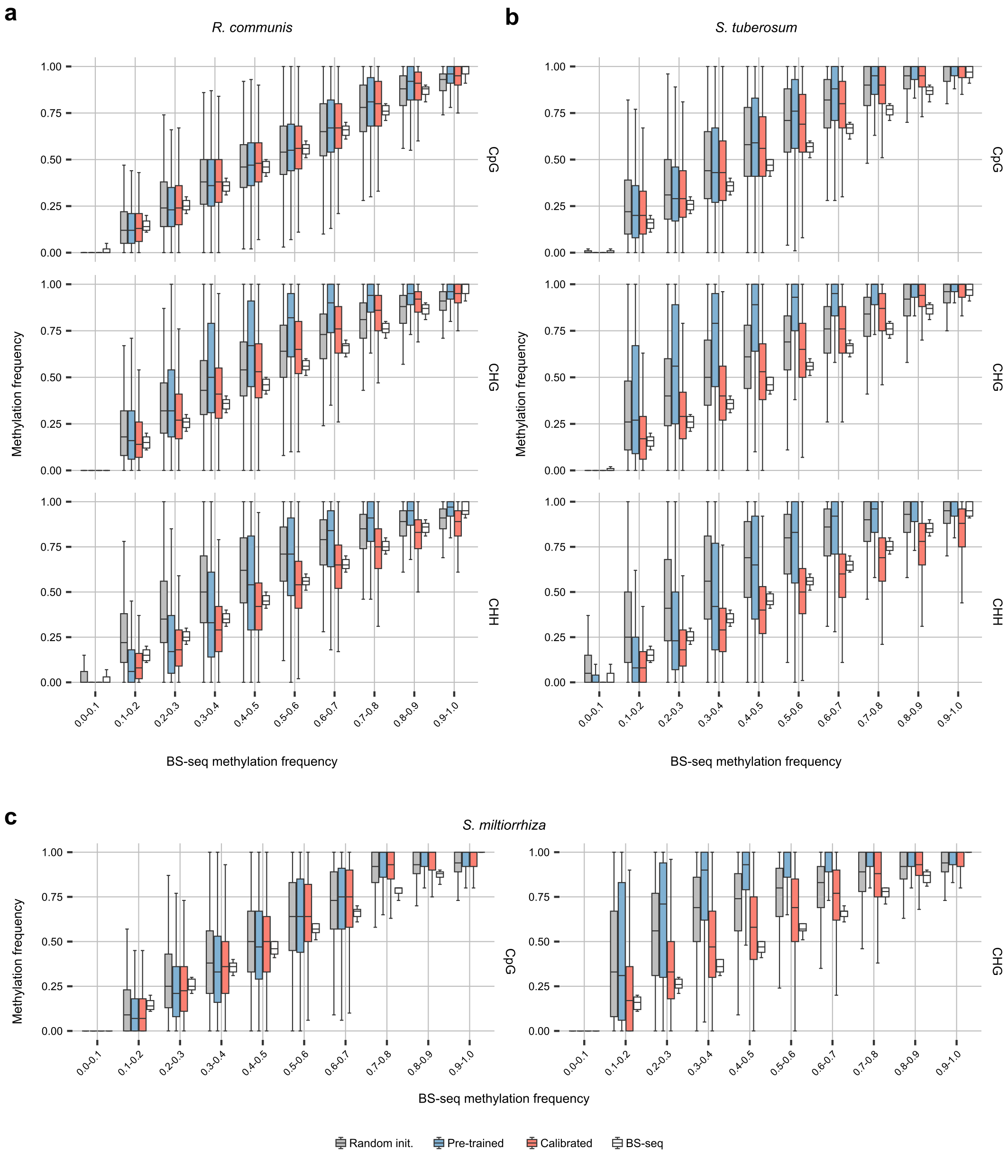

**Supplementary Fig. 4** Effects of pre-training and calibration on site-level methylation frequency estimates across BS-seq methylation bins. Site-level methylation frequencies predicted by the Random init., Pre-trained and Calibrated model variants and measured by BS-seq were stratified by BS-seq methylation frequency. **a,b** CpG, CHG and CHH methylation in *R. communis* (**a**) and *S. tuberosum* (**b**). **c** CpG and CHG methylation in *S. miltiorrhiza*; the corresponding CHH analysis is shown in Fig. 1c. Boxes show interquartile ranges, centre lines indicate medians, and whiskers extend to the most extreme values within 1.5 times the interquartile range; outliers are not shown.

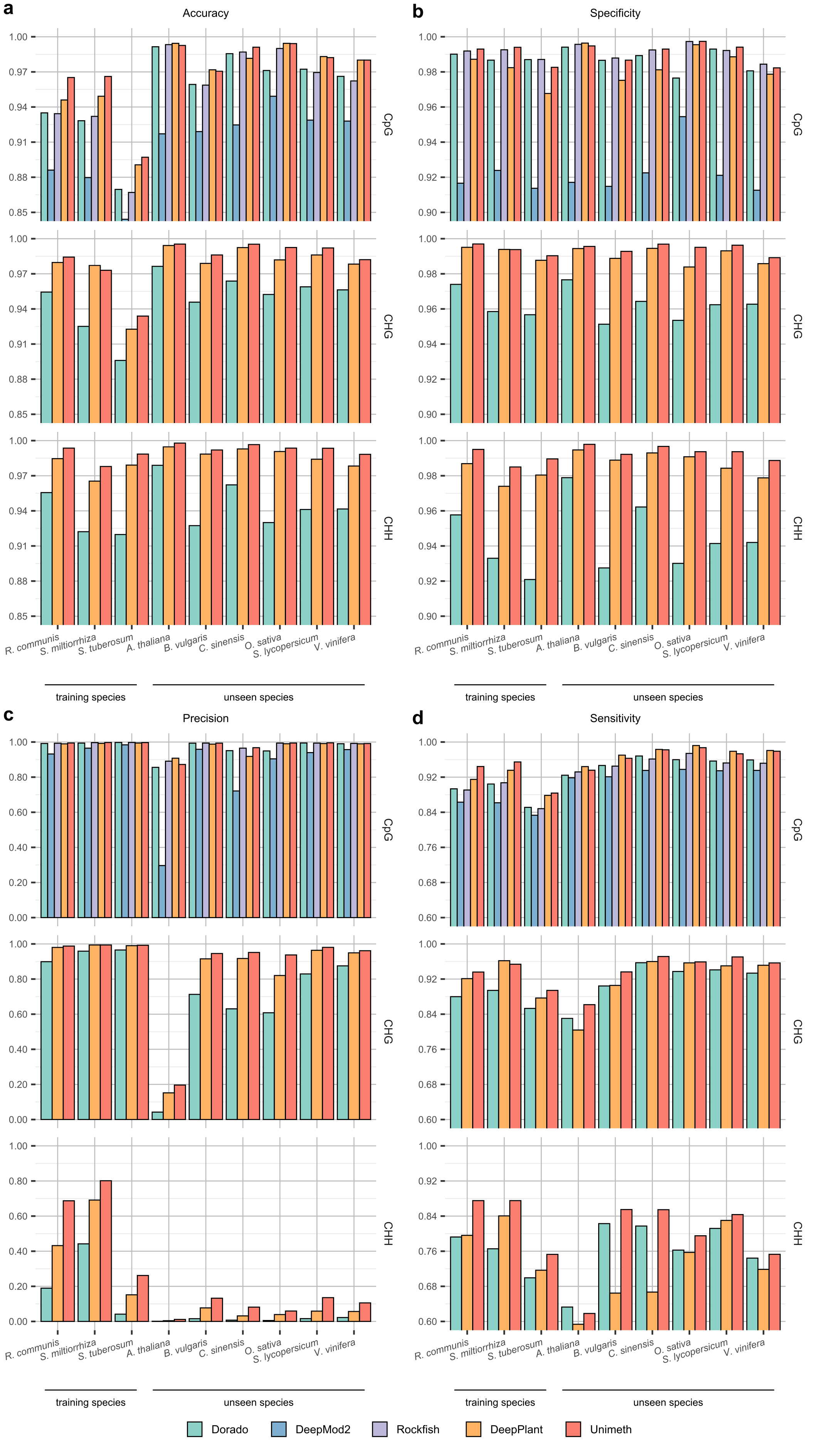

**Supplementary Fig. 5** Read-level evaluation of Unimeth and other methods on 5mC detection of plants using **nanopore R10.4.1 5kHz reads from Chen *et al***.^1^. **a** Accuracy. **b** Specificity. **c** Precision. **d** Sensitivity.

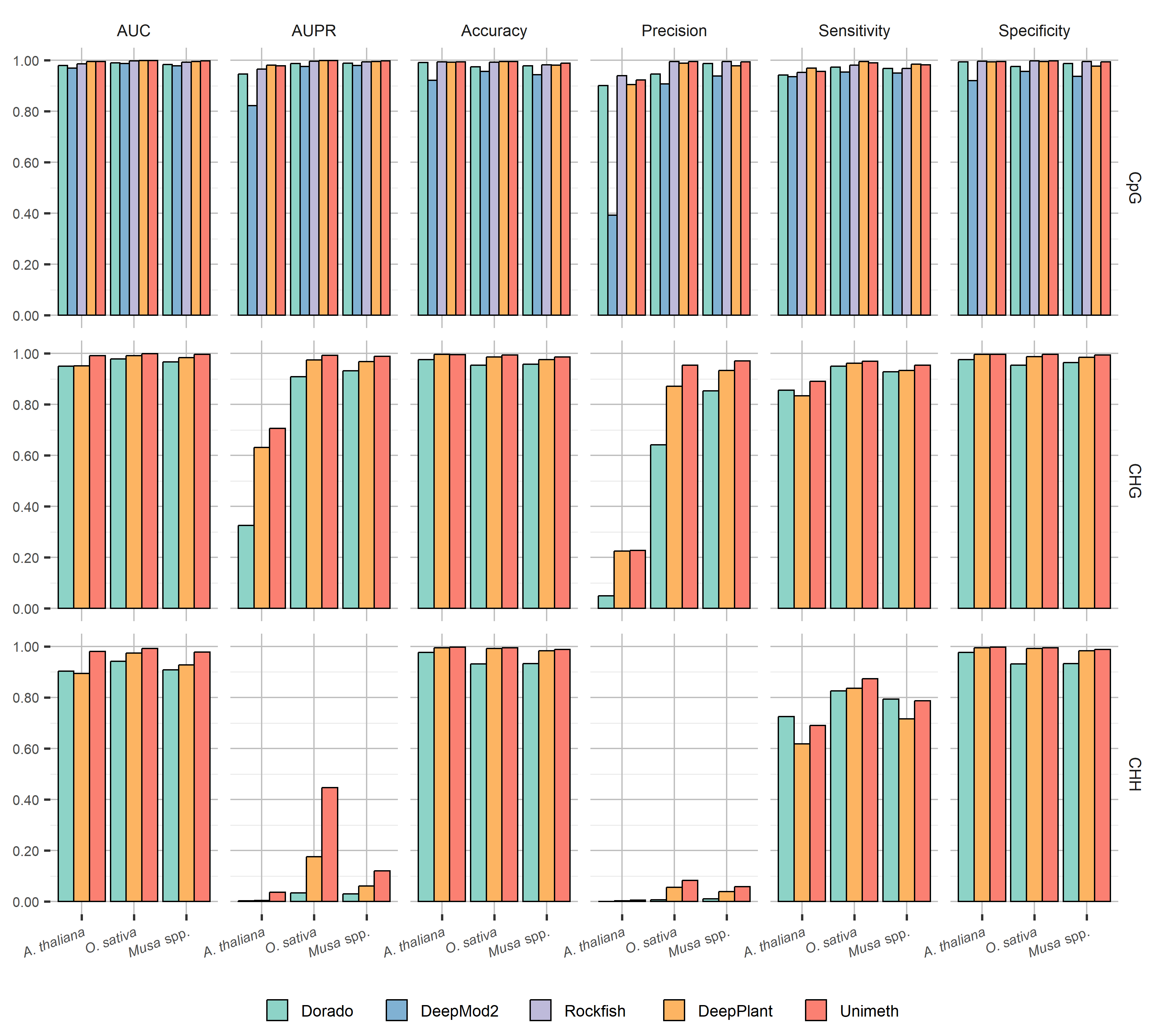

**Supplementary Fig. 6** Read-level evaluation of Unimeth and other methods on 5mC detection of plants using **in-house nanopore R10.4.1 5kHz reads**.

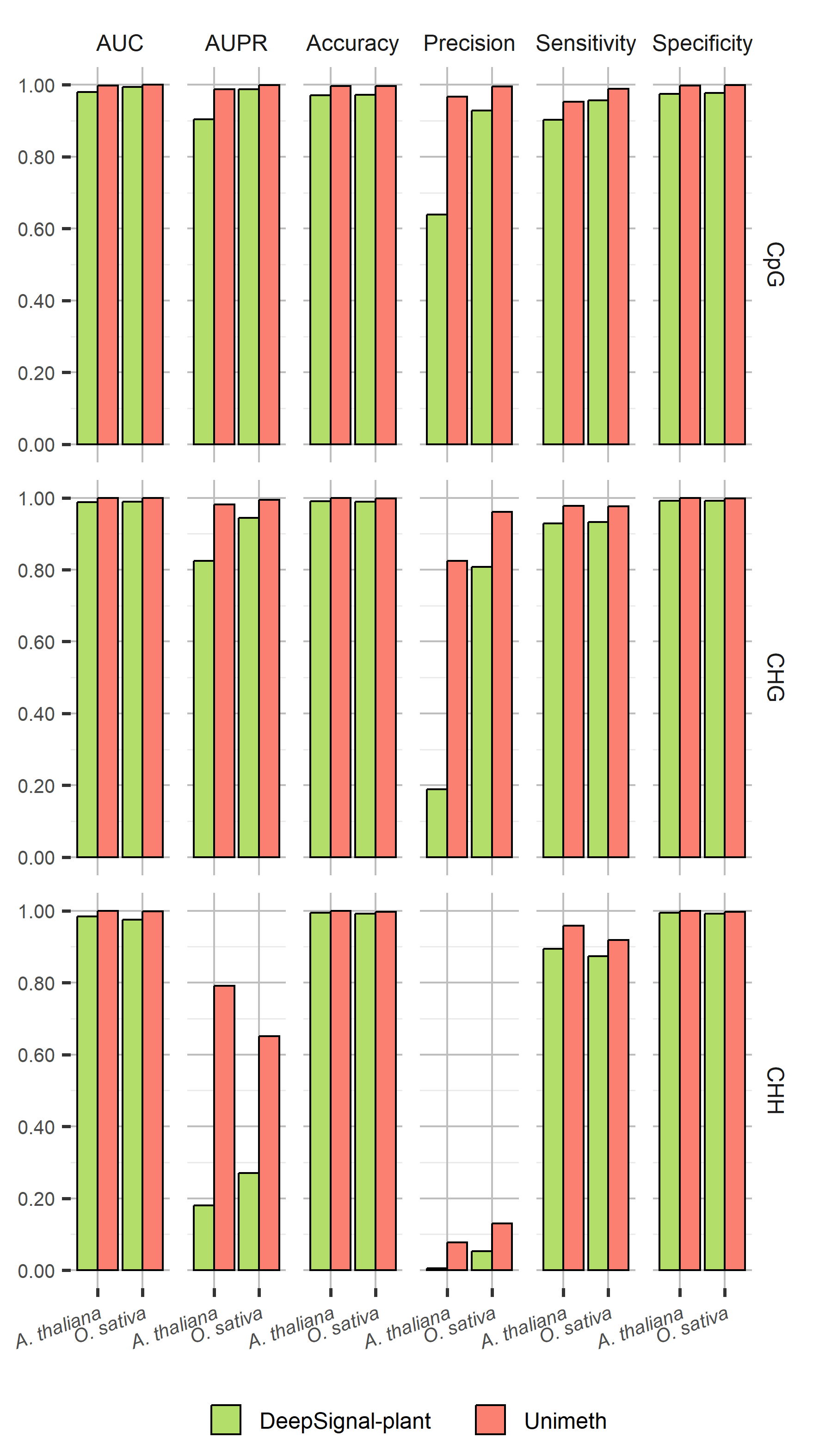

**Supplementary Fig. 7** Read-level evaluation of Unimeth and other methods on 5mC detection of plants using **nanopore R9.4.1 reads**.

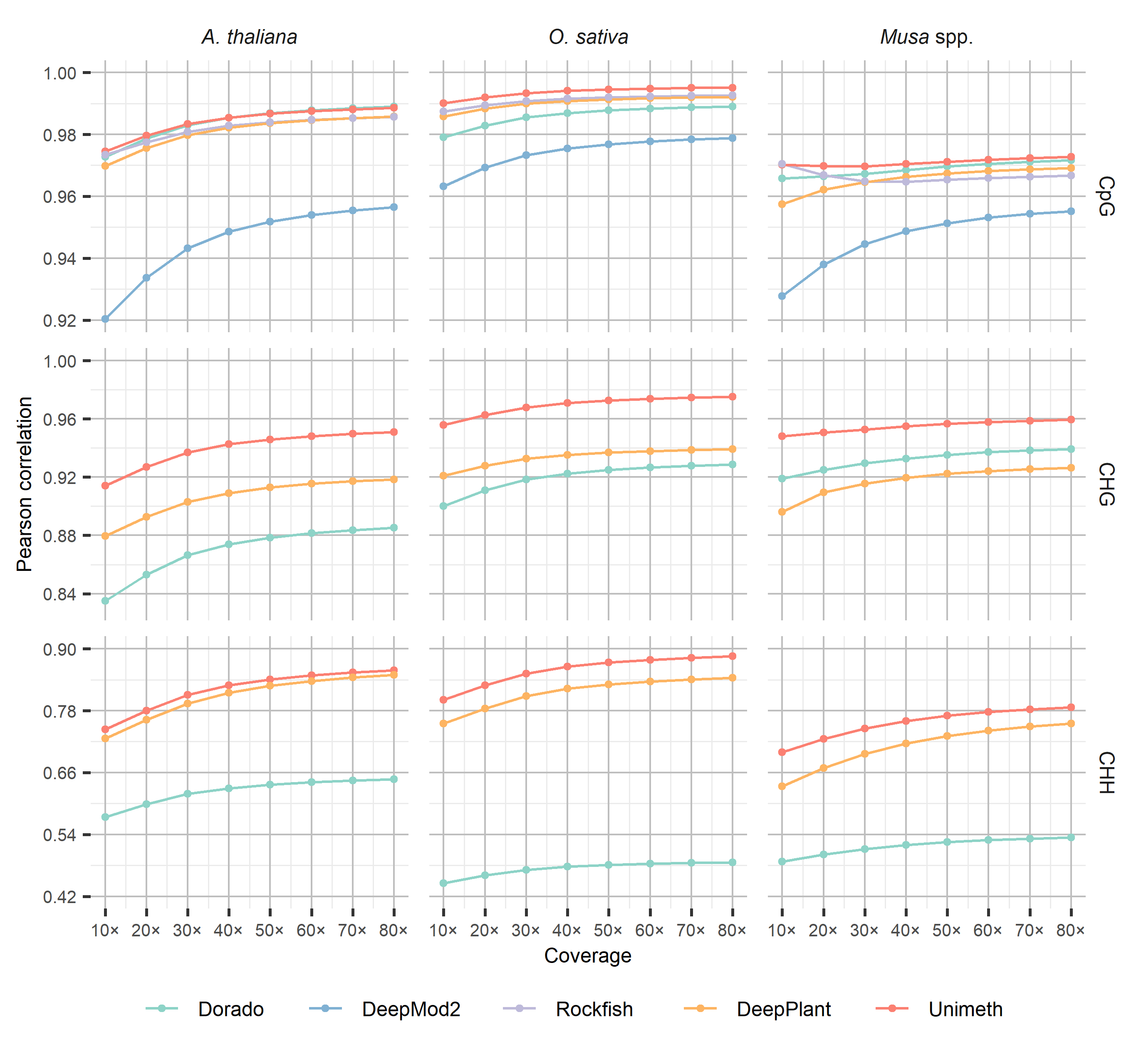

**Supplementary Fig. 8** Site-level evaluation of Unimeth and other methods on 5mC detection of plants using **in-house nanopore R10.4.1 5kHz reads**. Pearson correlation coefficients with BS-seq are shown across read coverages from 10× to 80× for CpG, CHG, and CHH contexts. Values represent the mean of five subsampling replicates; standard deviation values are provided in Supplementary Data 5.

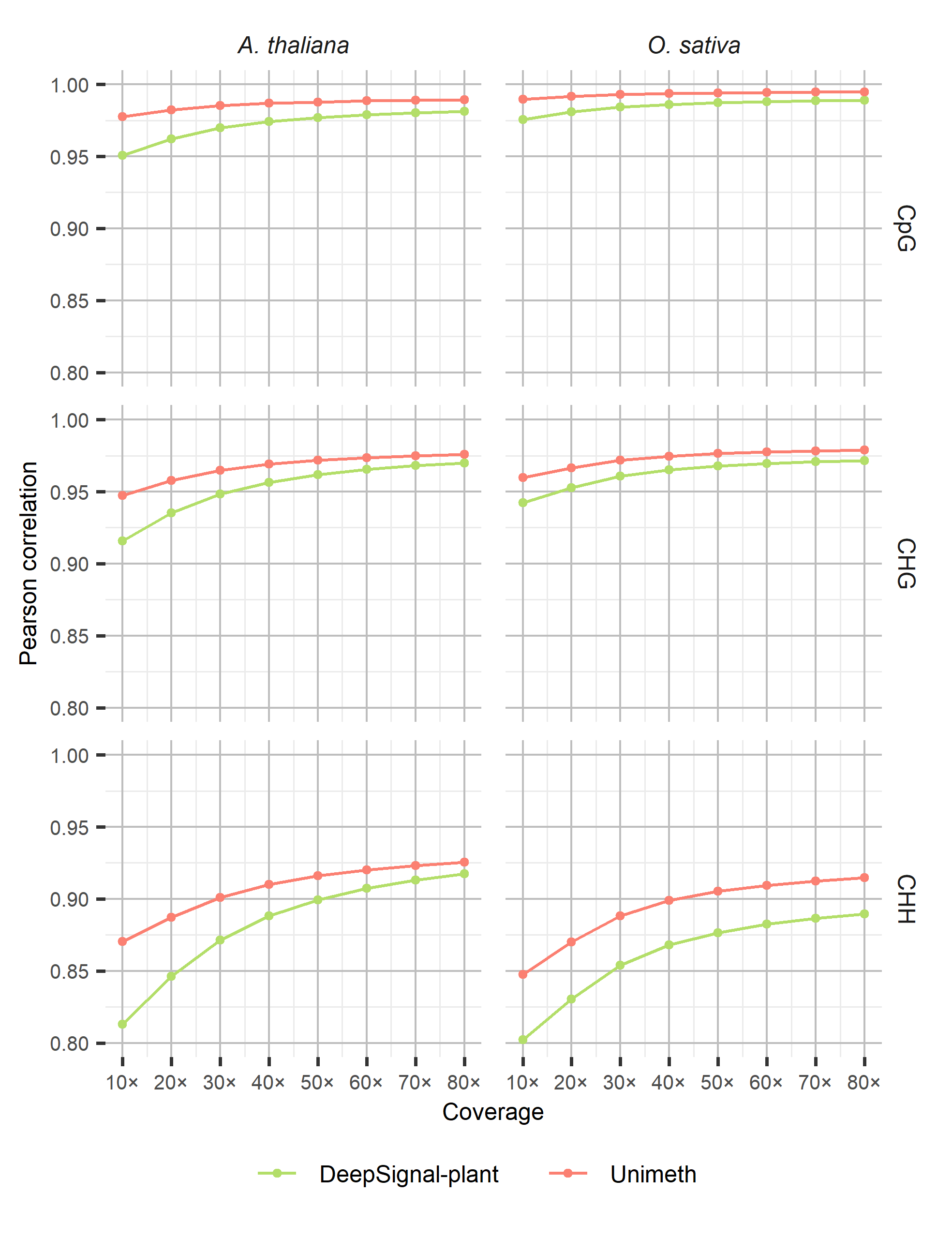

**Supplementary Fig. 9** Site-level evaluation of Unimeth and other methods on 5mC detection of plants using **nanopore R9.4.1 reads**. Pearson correlation coefficients with BS-seq are shown across read coverages from 10× to 80× for CpG, CHG, and CHH contexts. Values represent the mean of five subsampling replicates; standard deviation values are provided in Supplementary Data 6.

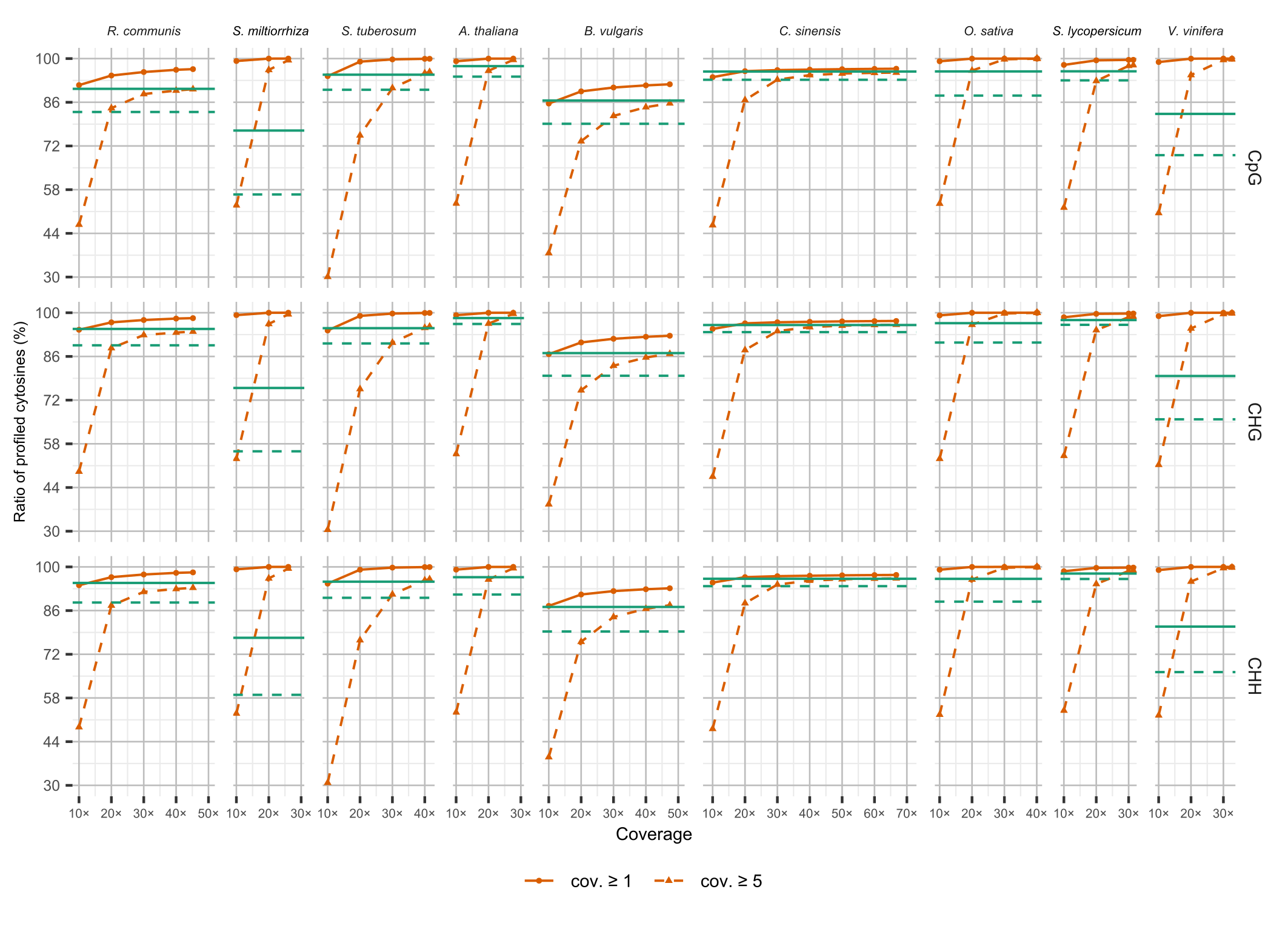

**Supplementary Fig. 10** The ratio of cytosines detected (*i.e.*, covered by at least 1 or 5 reads) under different coverages of **nanopore R10.4.1 5kHz reads from Chen *et al.***^1^ for CpG, CHG, and CHH motifs. For coverage values that are multiples of 10, values are the mean of five subsampling replicates. The solid and dashed horizontal lines in each panel indicate the fraction of cytosines covered with at least 1 and 5 reads by corresponding BS-seq data, respectively. For the nanopore data, the reads of the chromosomes/contigs for testing of corresponding species were used for the analysis (see Supplementary Data 2). For the BS-seq data, 64.52×, 71.50×, 23.15×, 65.23×, 68.47×, 128.23×, 41.46×, 68.29×, and 23.52× reads of the chromosomes/contigs for testing were analyzed for corresponding species, respectively. cov.: coverage.

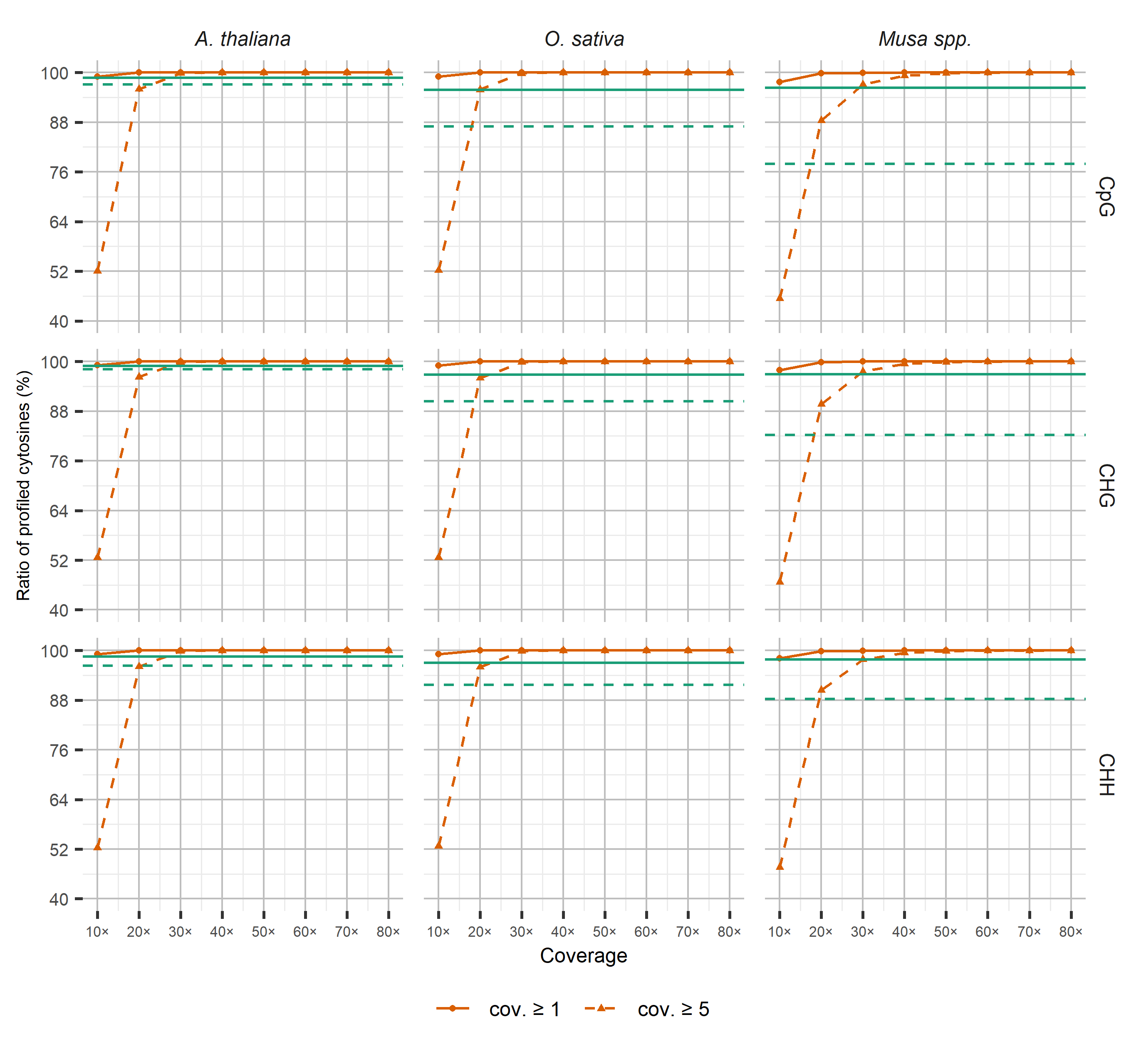

**Supplementary Fig. 11** The ratio of cytosines detected (*i.e.*, covered by at least 1 or 5 reads) under different coverages of **in-house nanopore R10.4.1 5kHz reads** for CpG, CHG, and CHH motifs. For coverage 10× to 80×, values are the mean of five subsampling replicates. The solid and dashed horizontal lines in each panel indicate the fraction of cytosines covered with at least 1 and 5 reads by corresponding BS-seq data, respectively. For the nanopore data, the reads of the chromosomes/contigs for testing of corresponding species were used for the analysis (see Supplementary Data 2). For the BS-seq data, 111.62×, 73.74×, and 35.93× reads of the chromosomes/contigs for testing were analyzed for corresponding species, respectively. cov.: coverage.

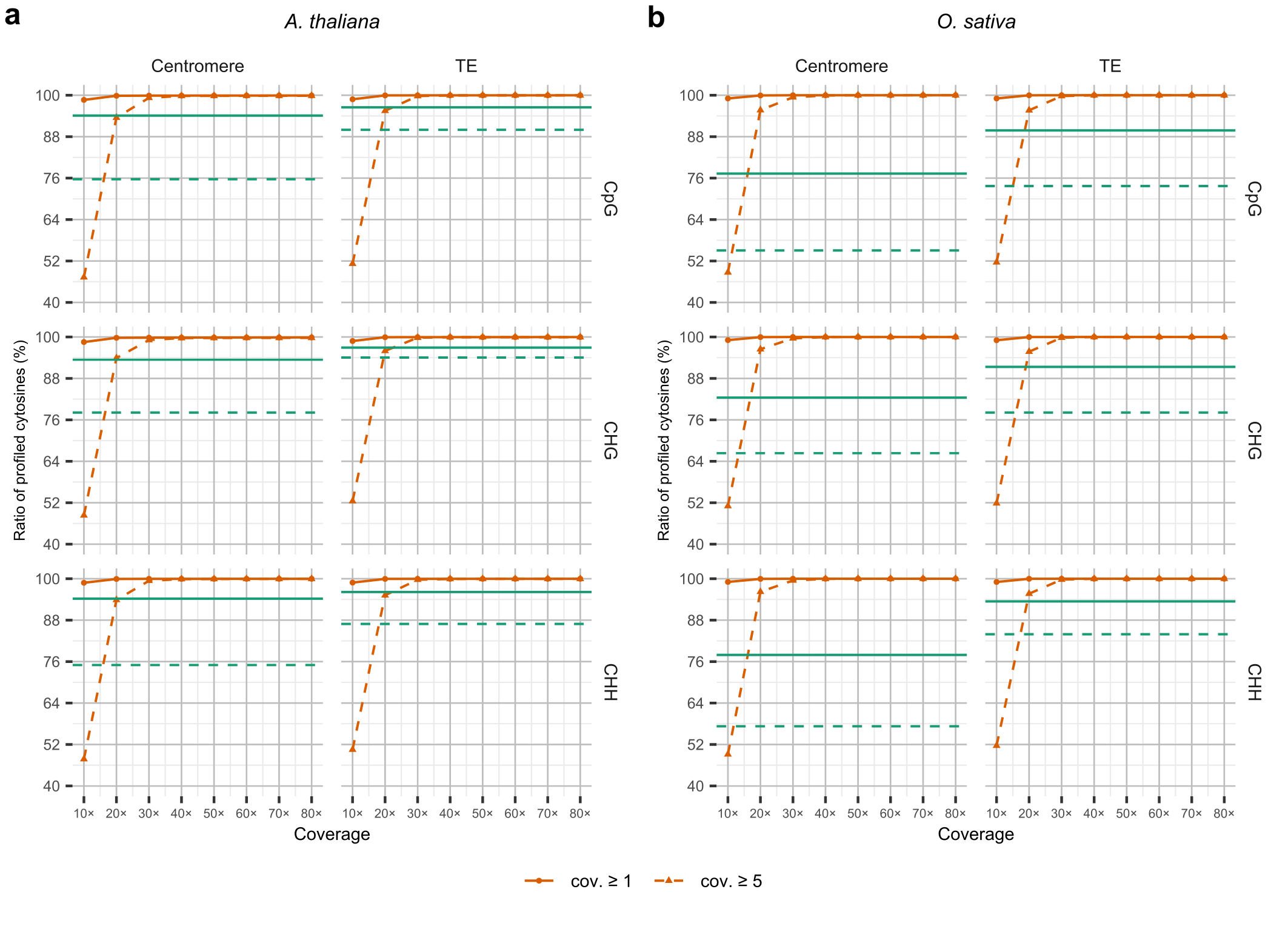

**Supplementary Fig. 12** The ratio of cytosines detected (*i.e.*, covered by at least 1 or 5 reads) in repetitive genomic regions under different coverages of nanopore reads. The **in-house nanopore R10.4.1 5kHz reads of chromosome 1 of *A. thaliana* and *O. sativa*** were used for the analysis. **a** *A. thaliana*. **b** *O. sativa*. For coverage 10×-80×, values are the mean of five subsampling replicates. The solid and dashed horizontal lines in each panel indicate the ratio of cytosines covered with at least 1 and 5 reads by corresponding BS-seq data, respectively. 111.62× and 73.74× reads of BS-seq for the chromosome 1 were analyzed for *A. thaliana* and *O. sativa*, respectively. cov.: coverage.

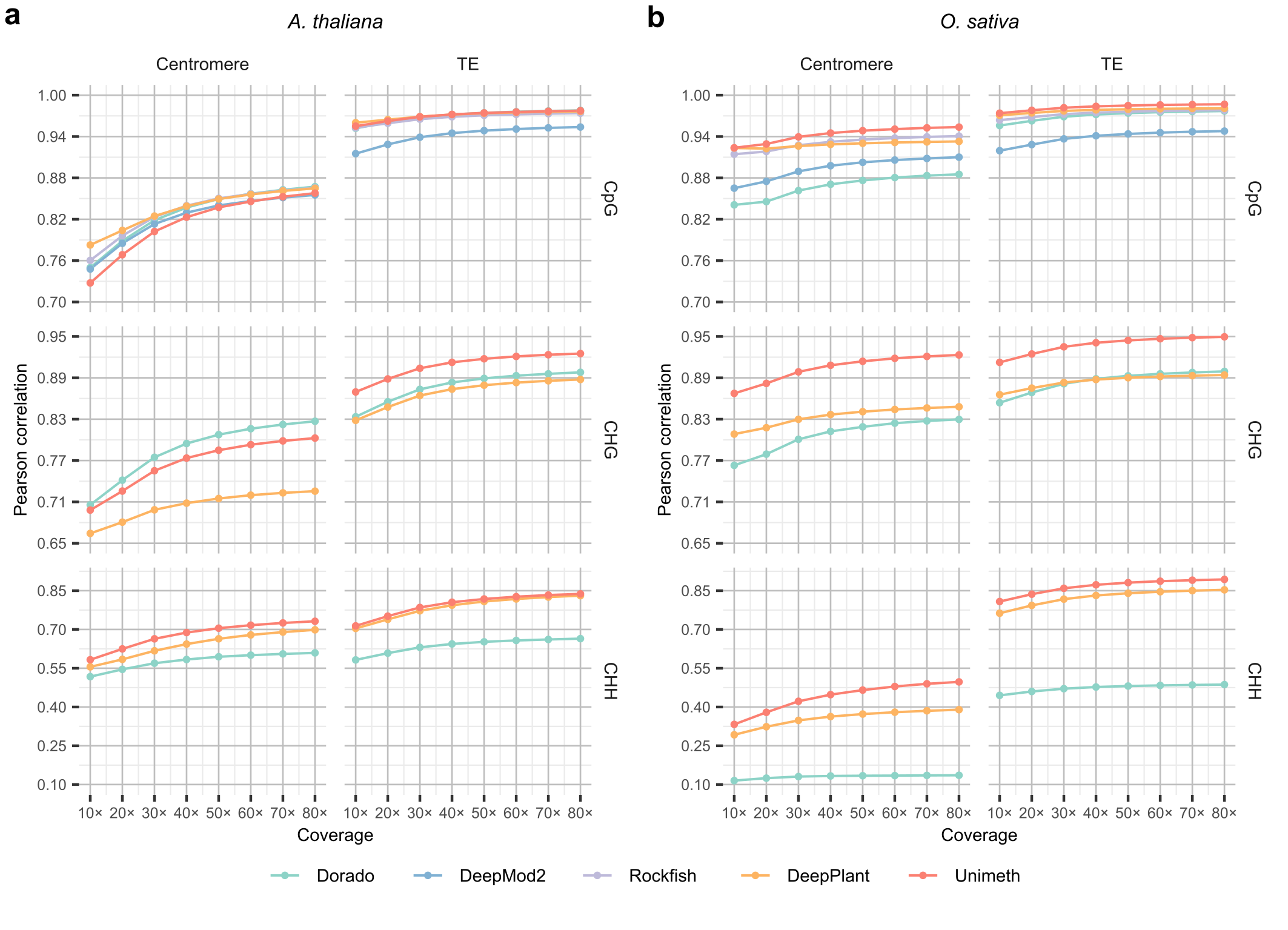

**Supplementary Fig. 13** Site-level evaluation of Unimeth and other methods on 5mC detection in repetitive genomic regions of plants using **in-house nanopore R10.4.1 5kHz reads**. **a** Pearson correlation with the results of BS-seq on *A. thaliana*. **b** Pearson correlation with the results of BS-seq on *O. sativa*. For coverage 10×-80×, values are the mean of five subsampling replicates. The standard deviation values of the multiple repeated tests are in Supplementary Data 7.

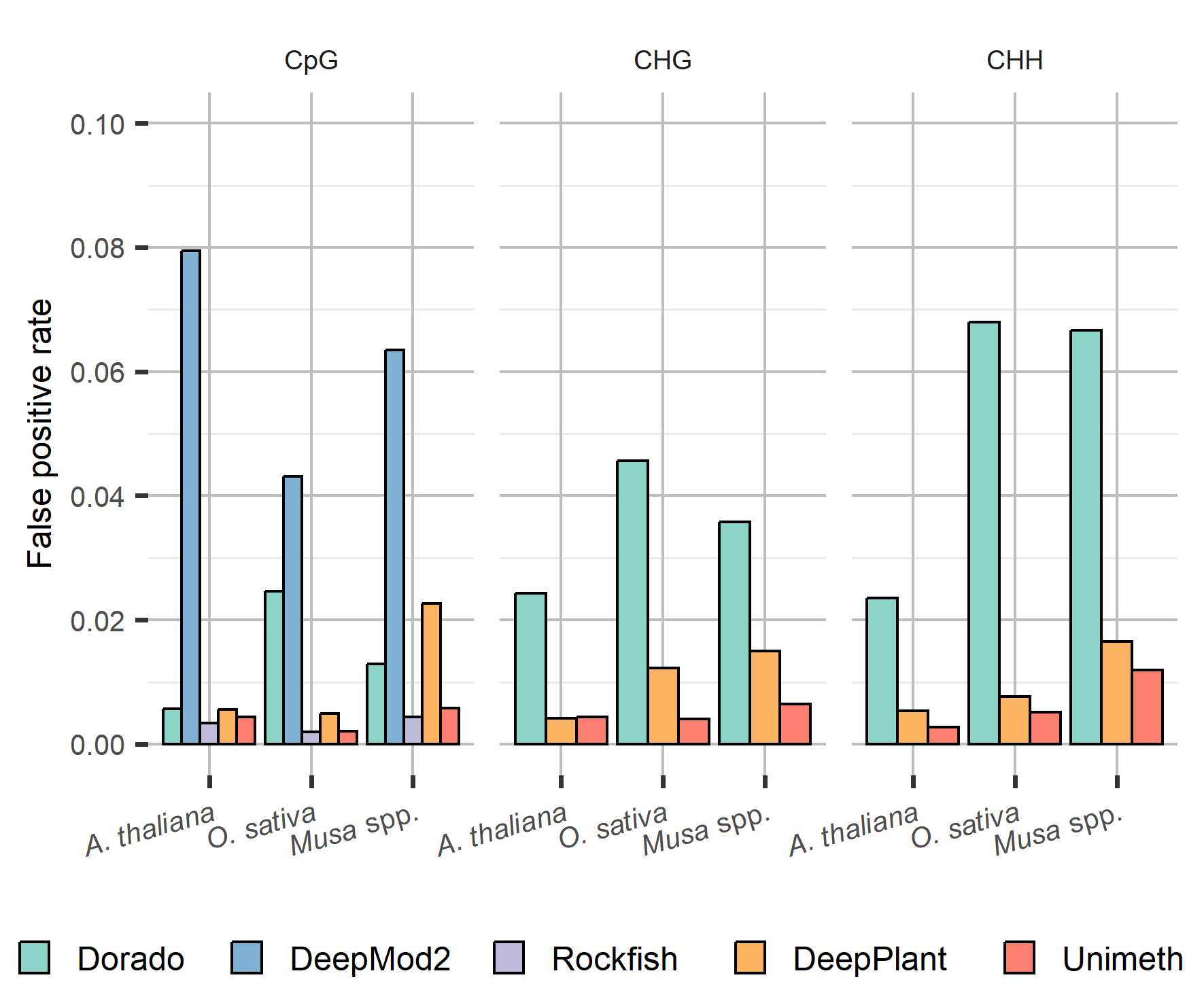

**Supplementary Fig. 14** Read-level false positive rates of Unimeth and other methods on 5mC detection of plants using **in-house nanopore R10.4.1 5kHz reads**.

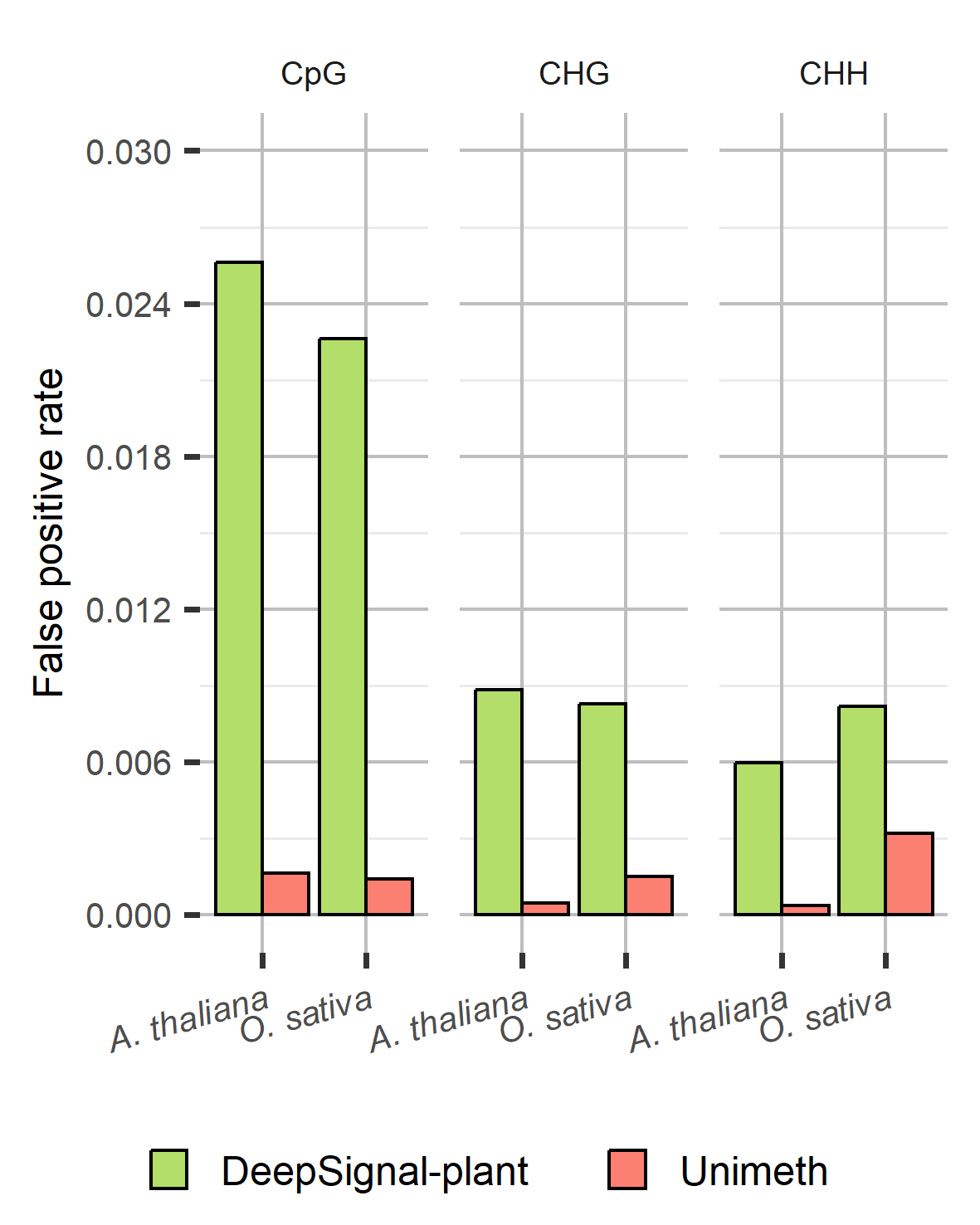

**Supplementary Fig. 15** Read-level false positive rates of Unimeth and other methods on 5mC detection of plants using **nanopore R9.4.1 reads**.

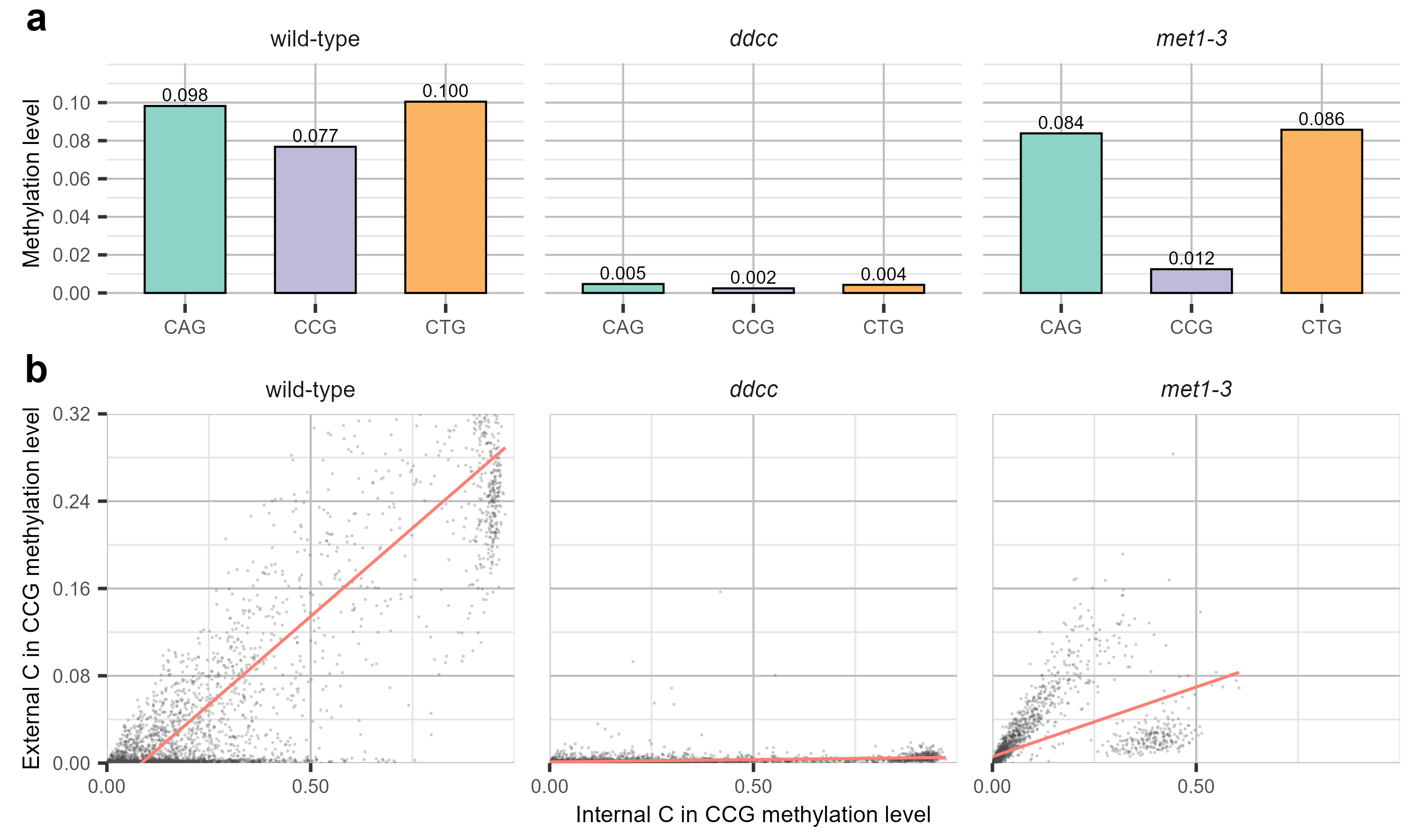

**Supplementary Fig. 16** CHG context-specific methylation and paired CCG methylation in *A. thaliana* mutants.

**a** Unimeth methylation levels at the three CHG sequence contexts, CAG, CCG and CTG, in *A. thaliana* Col-0 wild type, *ddcc* and *met1-3*. Values above bars indicate count-weighted methylation levels calculated as the ratio of methylated calls to total covered calls in 10-kb genomic windows. **b** Relationship between methylation of the external CHG-context cytosine and the internal CpG-context cytosine in CCG motifs. Each point represents a 10-kb genomic window; red lines indicate linear regression fits.

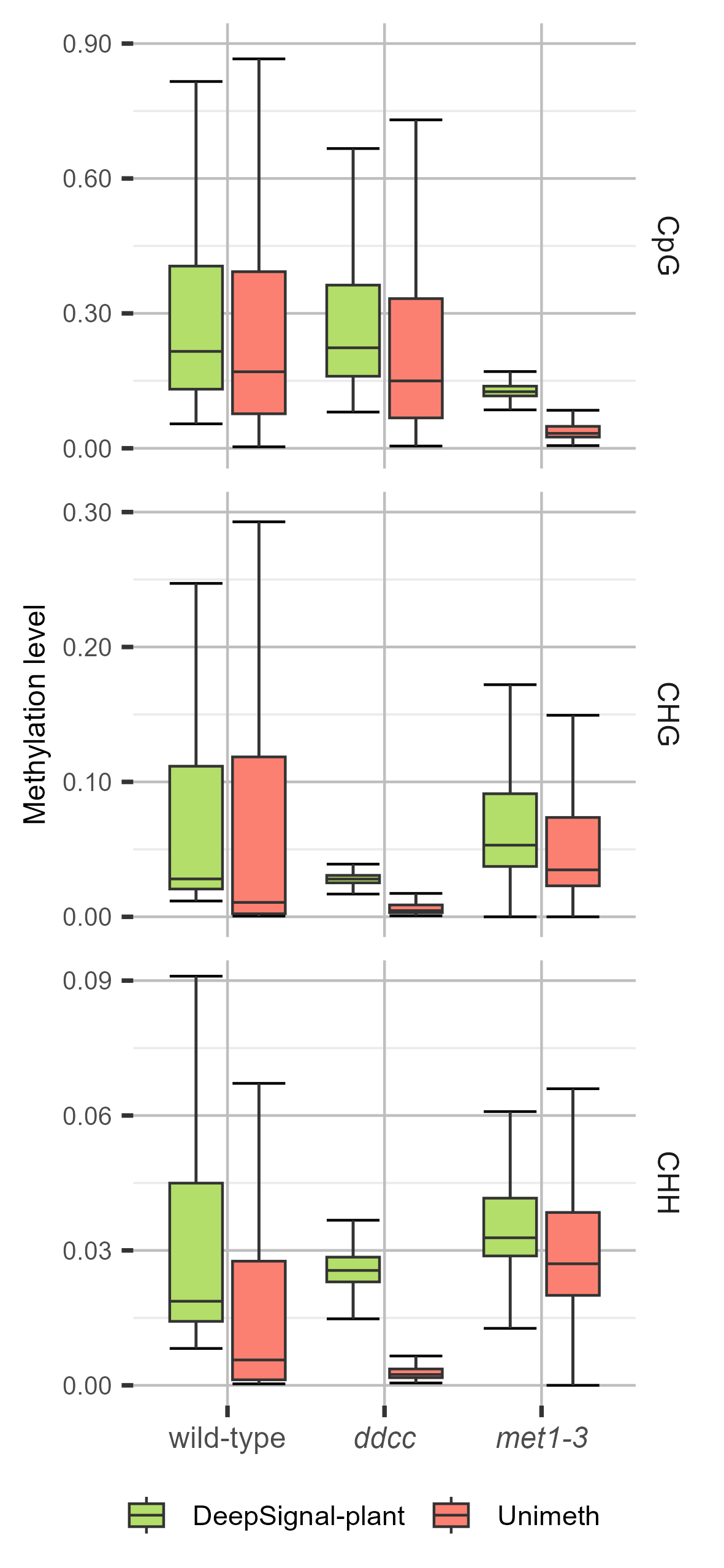

**Supplementary Fig. 17** Methylation levels detected by DeepSignal-plant and Unimeth in R9.4.1 nanopore data of *A. thaliana* Col-0 wild type and methylation-deficient mutants *ddcc* and *met1-3*. Methylation levels were calculated for 10-kb windows as the ratio of methylated calls to total covered calls within each window. Boxes show interquartile ranges, centre lines show medians, and whiskers indicate 1.5 times the interquartile range.

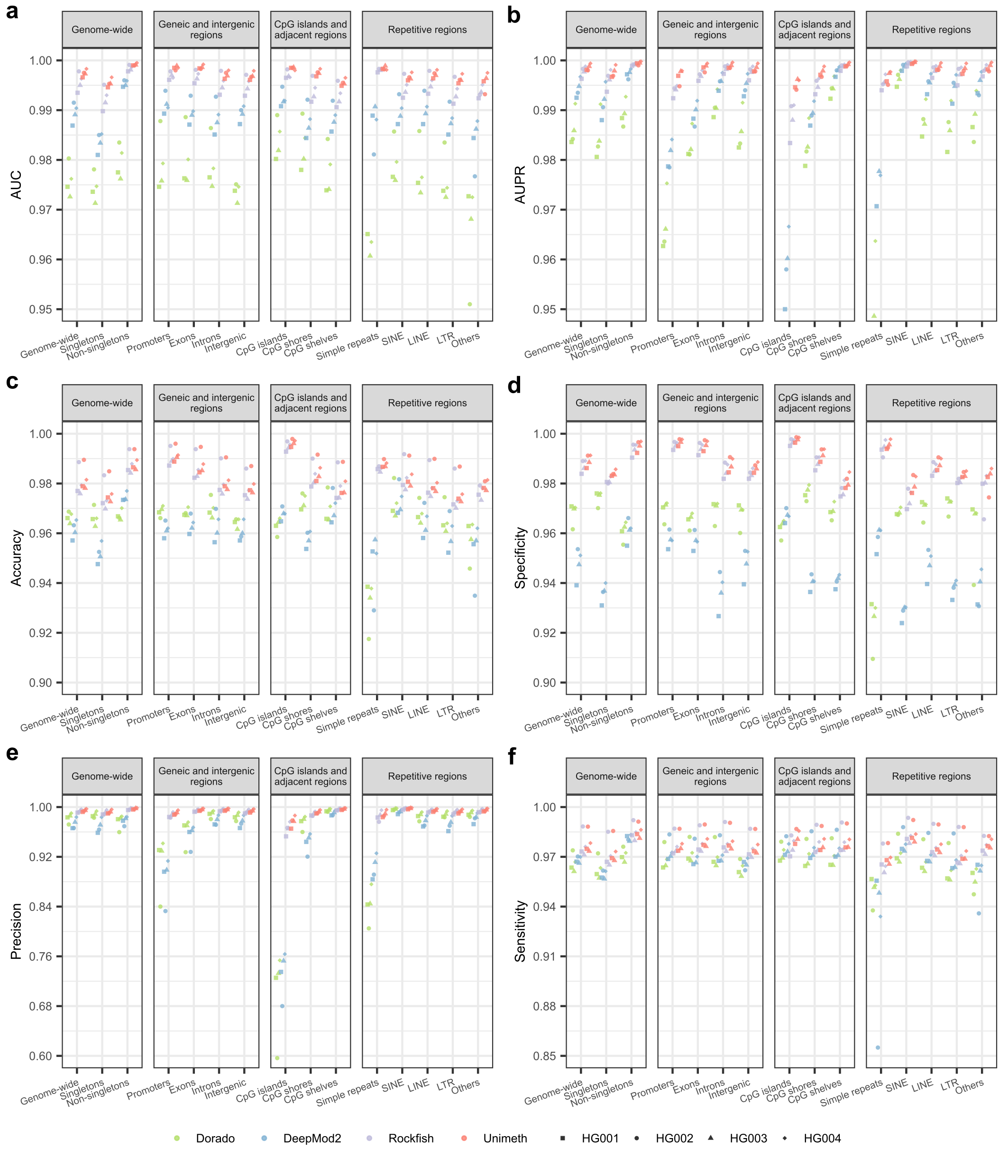

**Supplementary Fig. 18** Read-level evaluation of Unimeth and other methods on 5mCpG detection across different genomic contexts and regions in human (HG001-HG004) using **nanopore R10.4.1 5kHz reads**. **a** AUC. **b** AUPR. **c** Accuracy. **d** Specificity. **e** Precision. **f** Sensitivity.

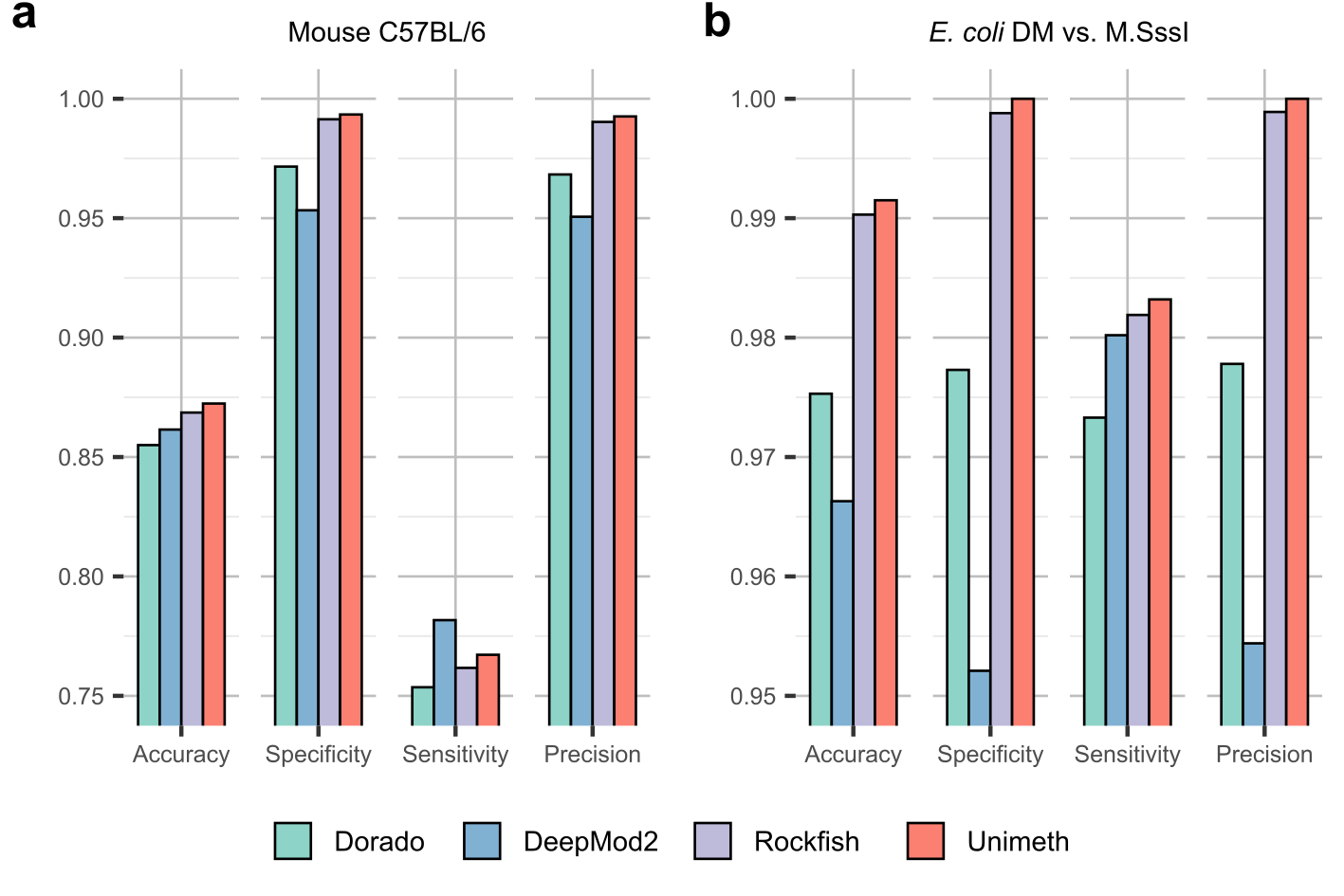

**Supplementary Fig. 19** Read-level evaluation of Unimeth and other methods on 5mCpG detection of mouse (**a**) and *E. coli* (**b**) using **nanopore R10.4.1 5kHz reads**.

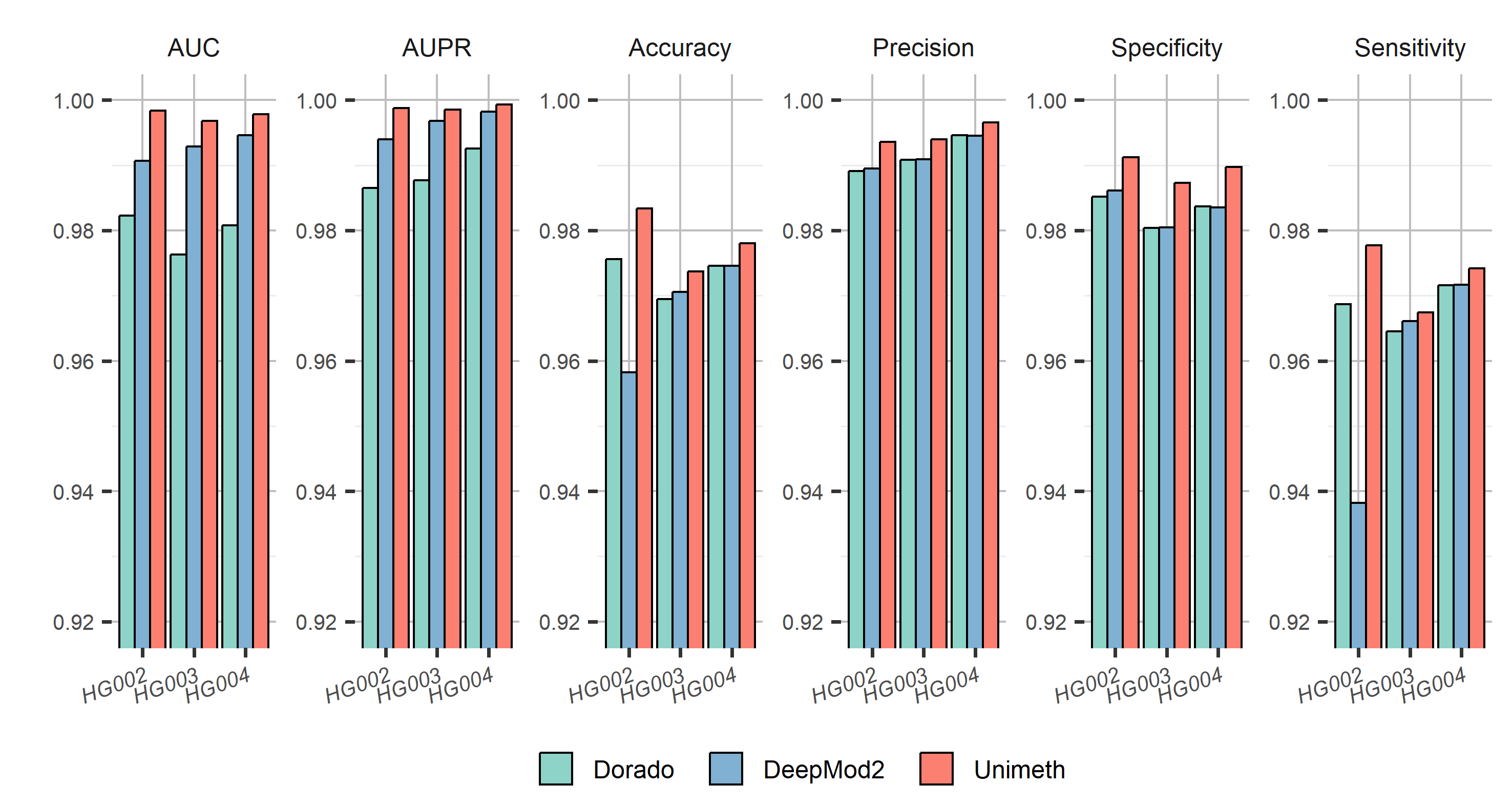

**Supplementary Fig. 20** Read-level evaluation of Unimeth and other methods on 5mCpG detection of human using **nanopore R10.4.1 4kHz reads**.

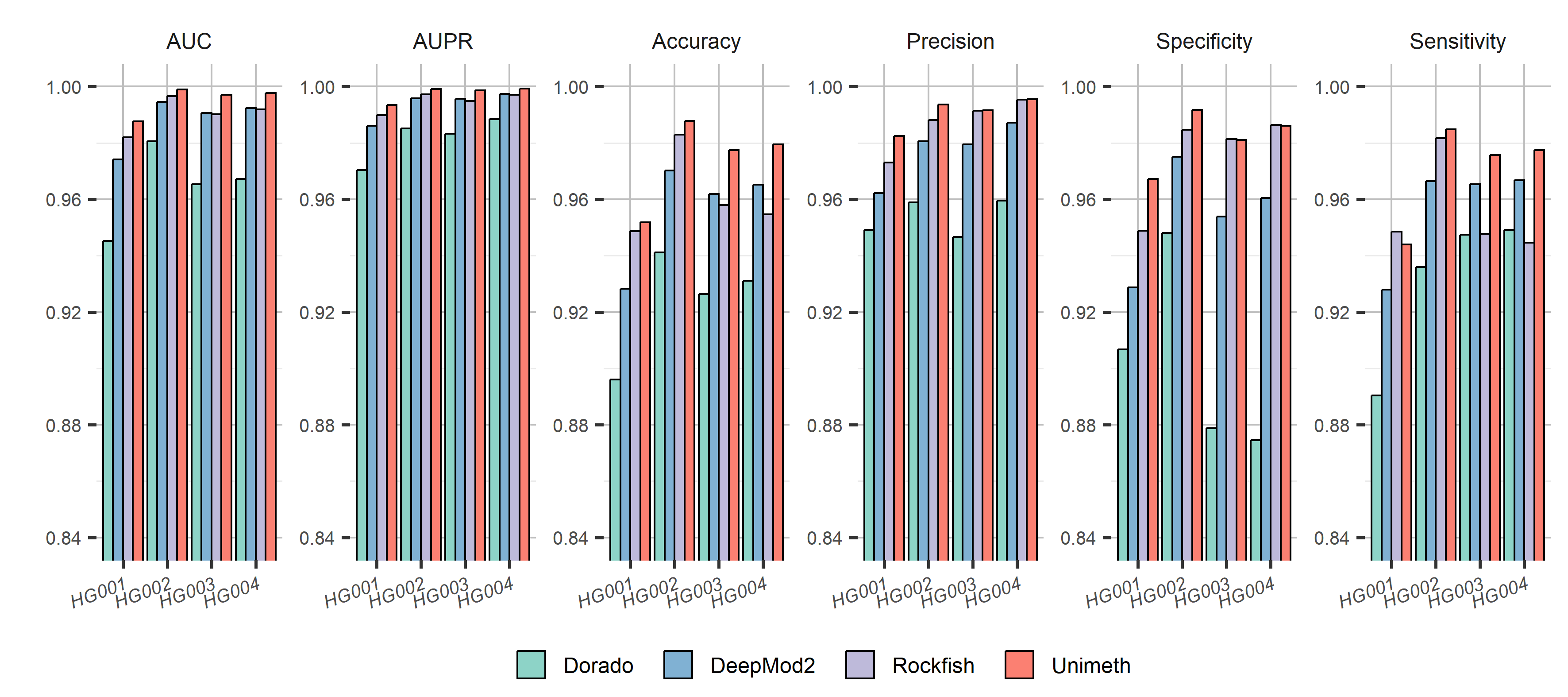

**Supplementary Fig. 21** Read-level evaluation of Unimeth and other methods on 5mCpG detection of human using **nanopore R9.4.1 reads**.

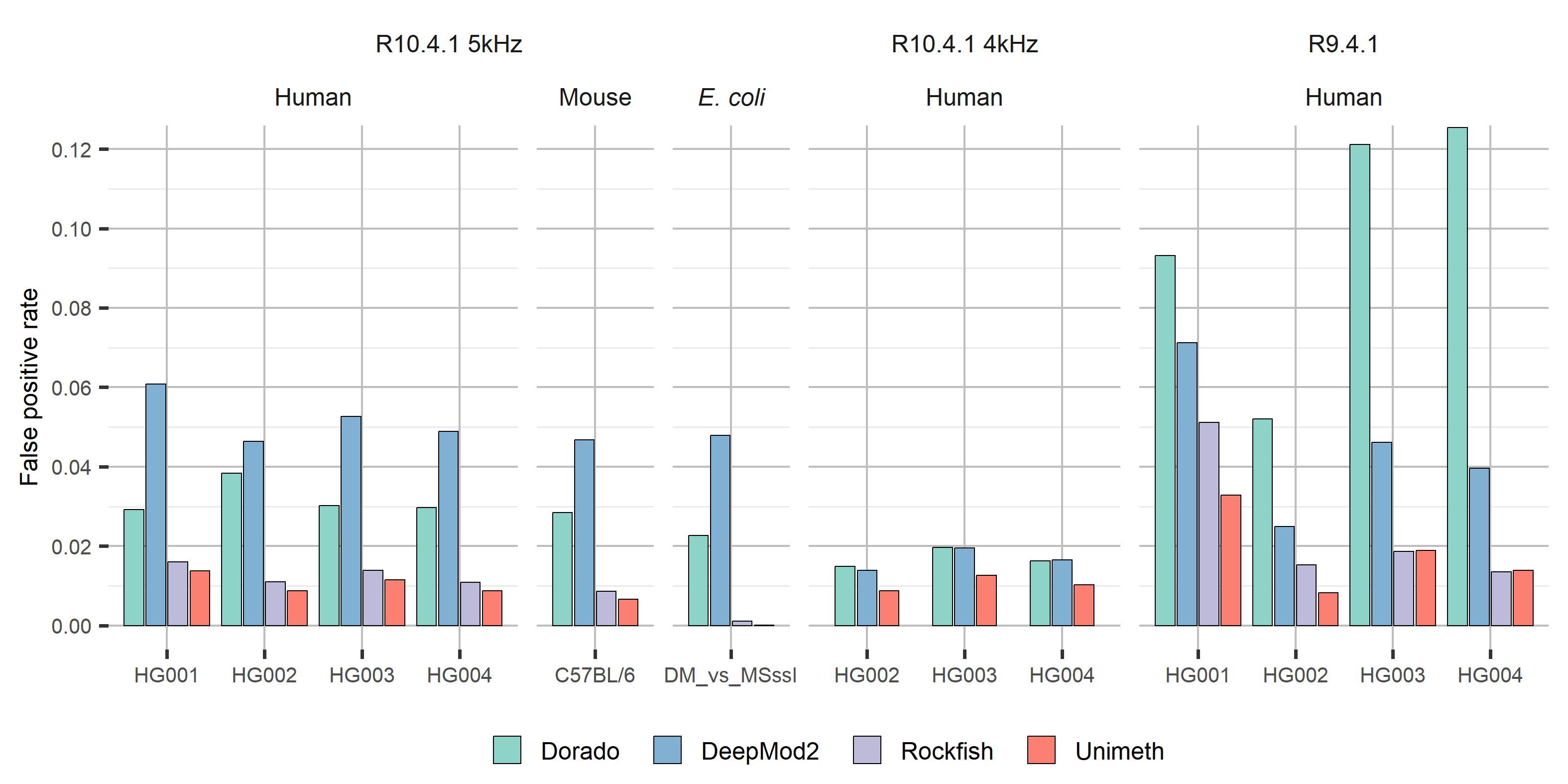

**Supplementary Fig. 22** Read-level false positive rates of Unimeth and other methods on 5mCpG detection.

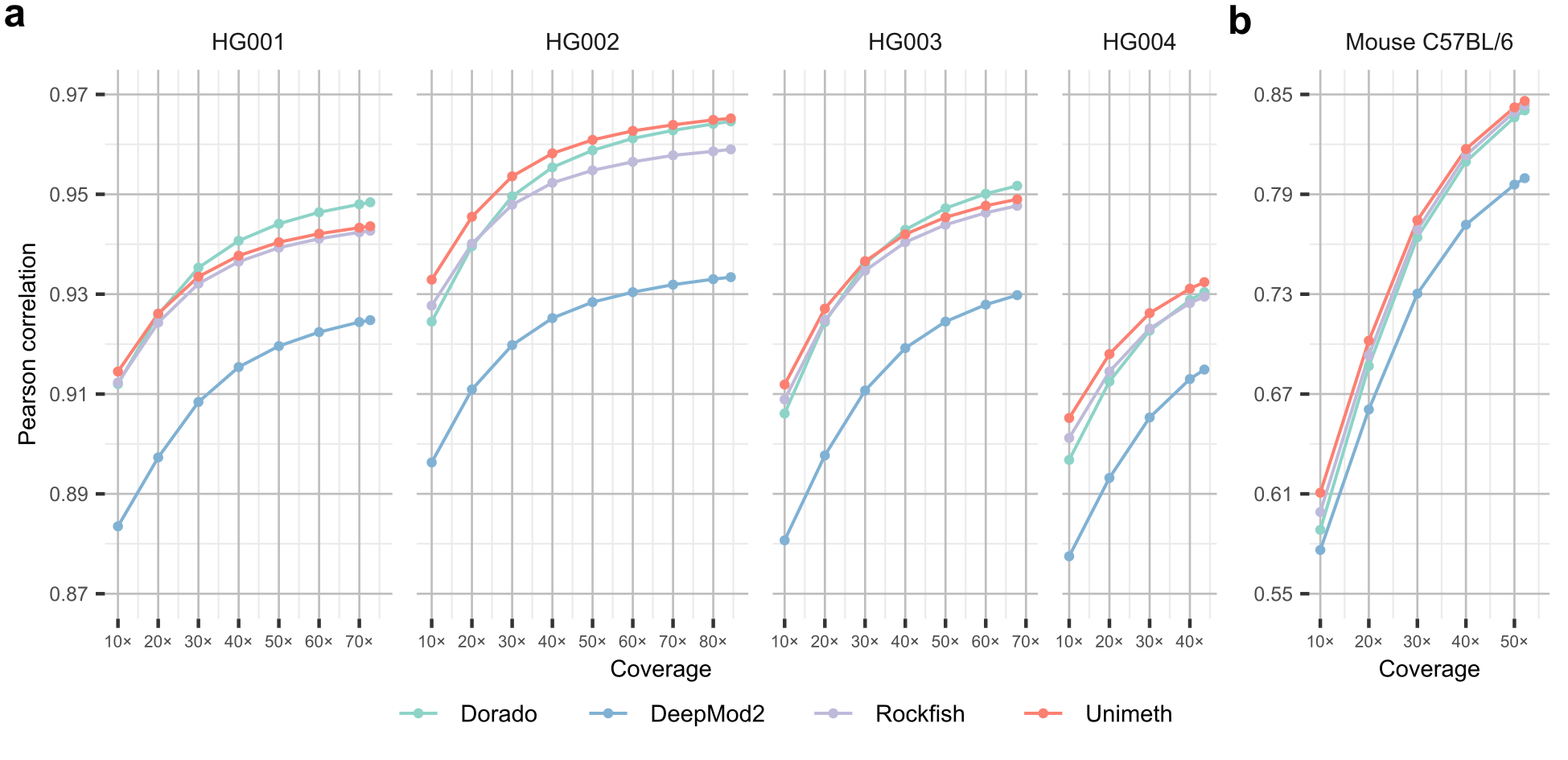

**Supplementary Fig. 23** Site-level evaluation of Unimeth and other methods on 5mCpG detection of human and mouse using **nanopore R10.4.1 5kHz reads**. **a** Human HG001-HG004. **b** Mouse C57BL/6. Values at subsampled coverages represent the mean of five subsampling replicates; standard deviation values are provided in Supplementary Data 8.

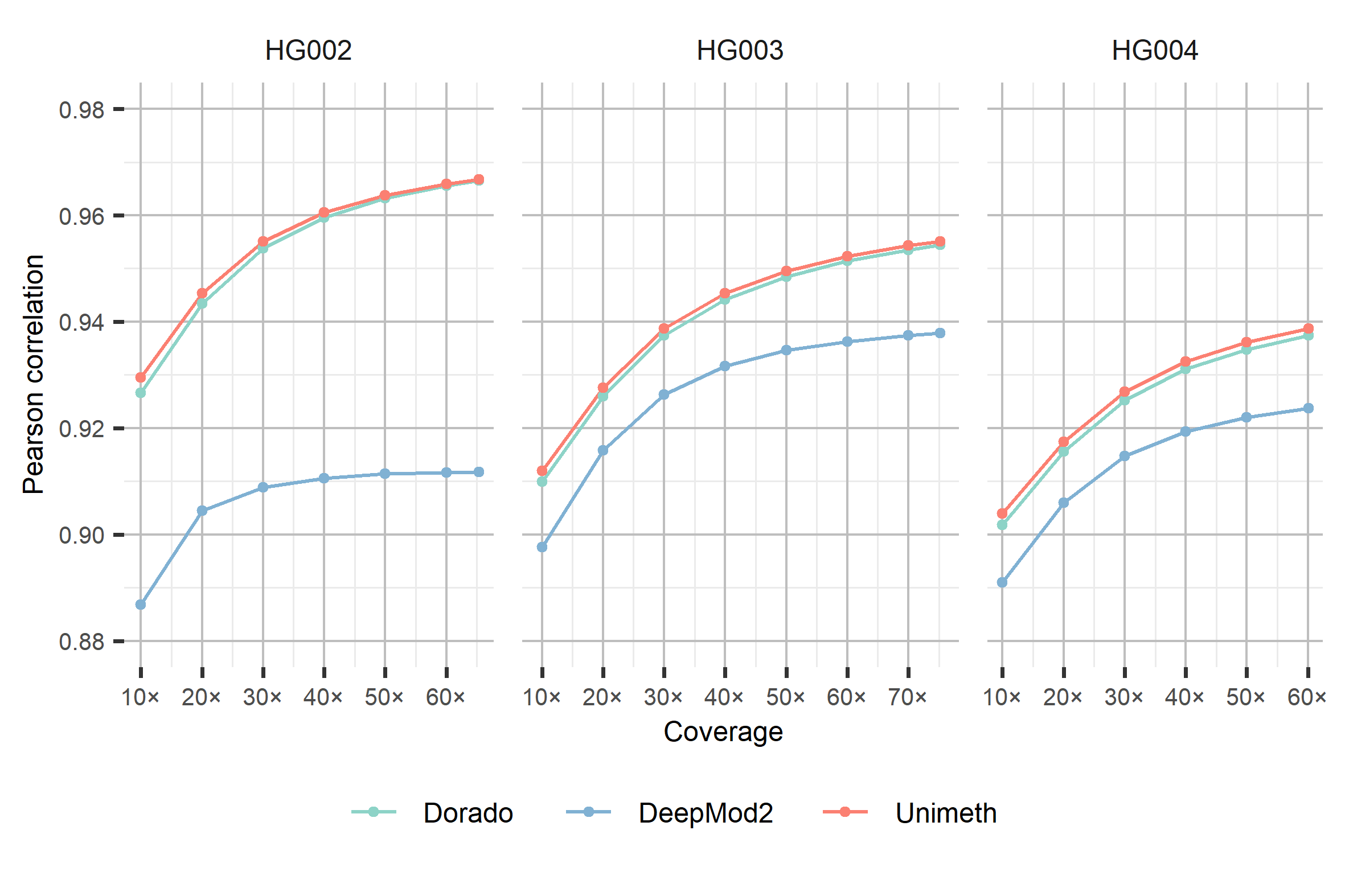

**Supplementary Fig. 24** Site-level evaluation of Unimeth and other methods on 5mCpG detection of human using **nanopore R10.4.1 4kHz reads**. Values at subsampled coverages represent the mean of five subsampling replicates; standard deviation values are provided in Supplementary Data 9.

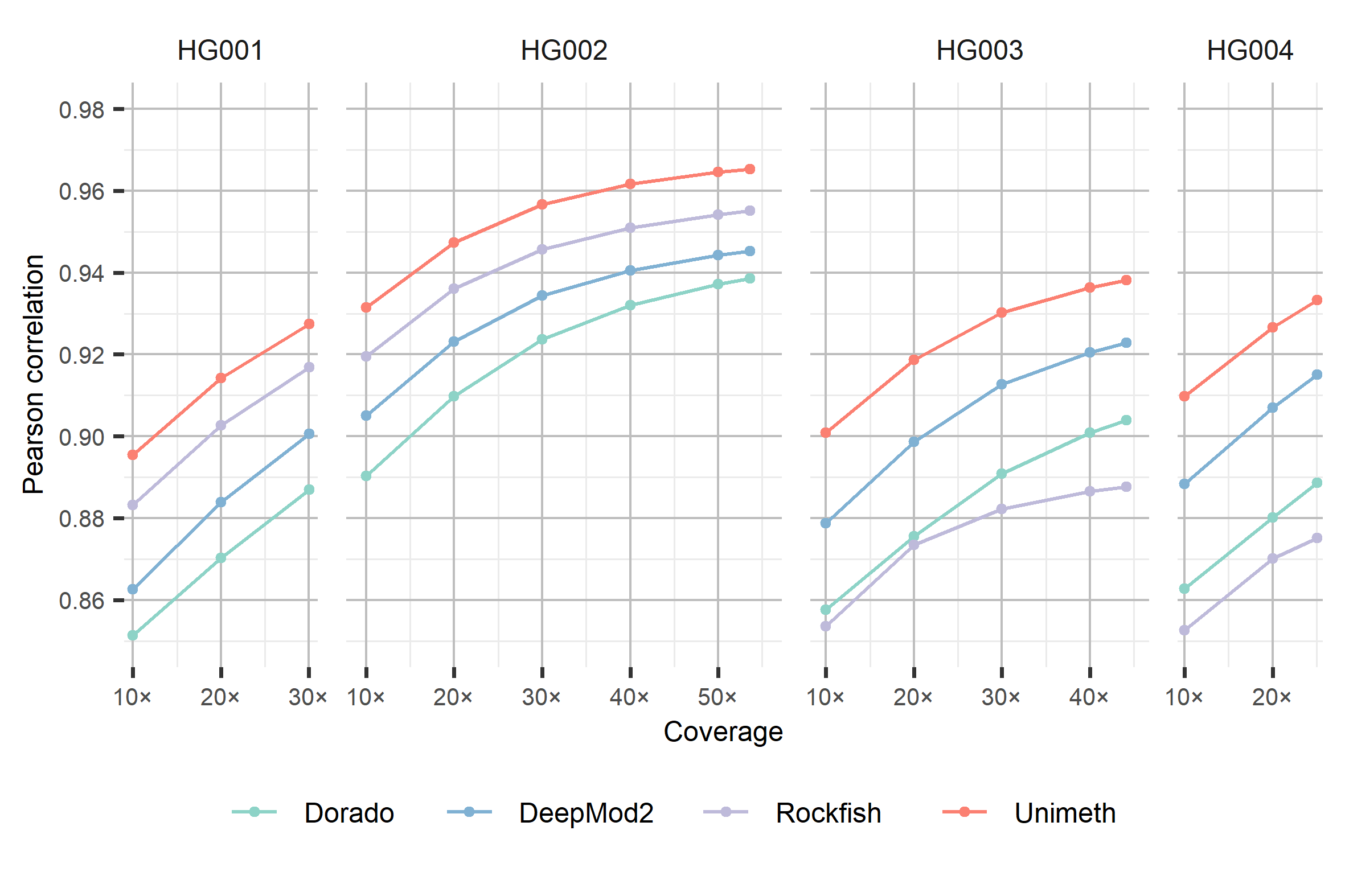

**Supplementary Fig. 25** Site-level evaluation of Unimeth and other methods on 5mCpG detection of human using **nanopore R9.4.1 reads**. Values at subsampled coverages represent the mean of five subsampling replicates; standard deviation values are provided in Supplementary Data 10.

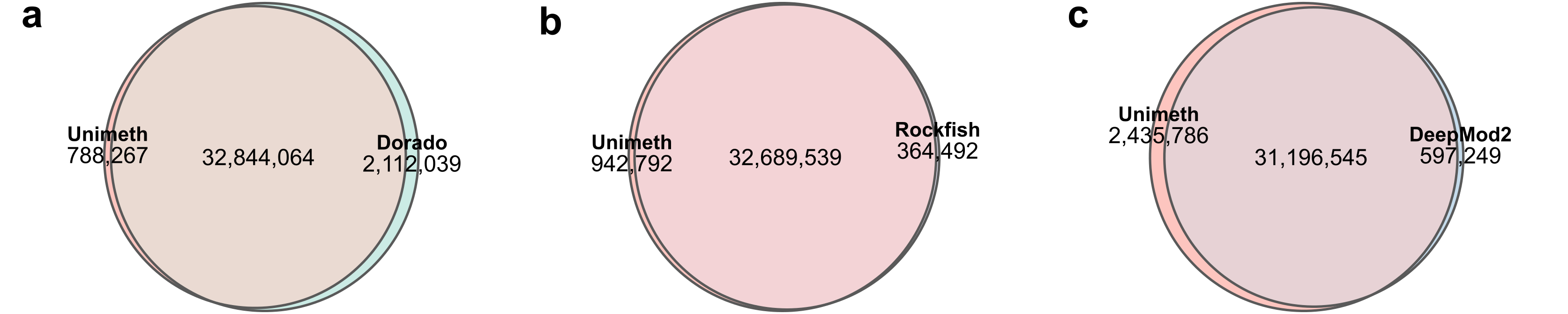

**Supplementary Fig. 26** Overlap of highly methylated CpG sites between Unimeth and other methods in HBA20-2. **a-c** Pairwise comparisons with Dorado (**a**), Rockfish (**b**) and DeepMod2 (**c**).

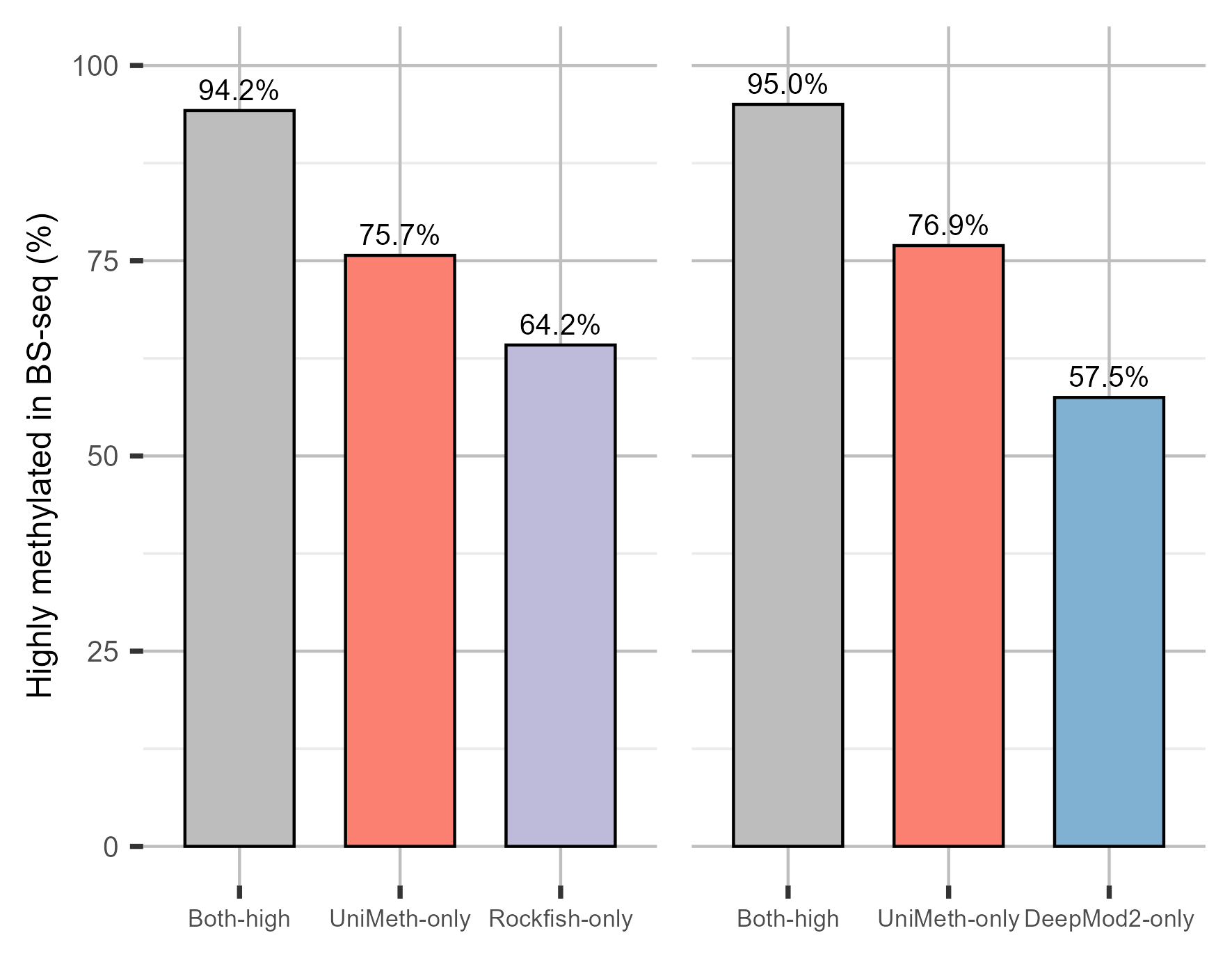

**Supplementary Fig. 27** Comparison of highly methylated CpG sites in HBA20-2. Unimeth is compared with Rockfish (left) and DeepMod2 (right). Bars show the percentages of sites also classified as highly methylated by BS-seq among sites called highly methylated by both methods or by either method alone.

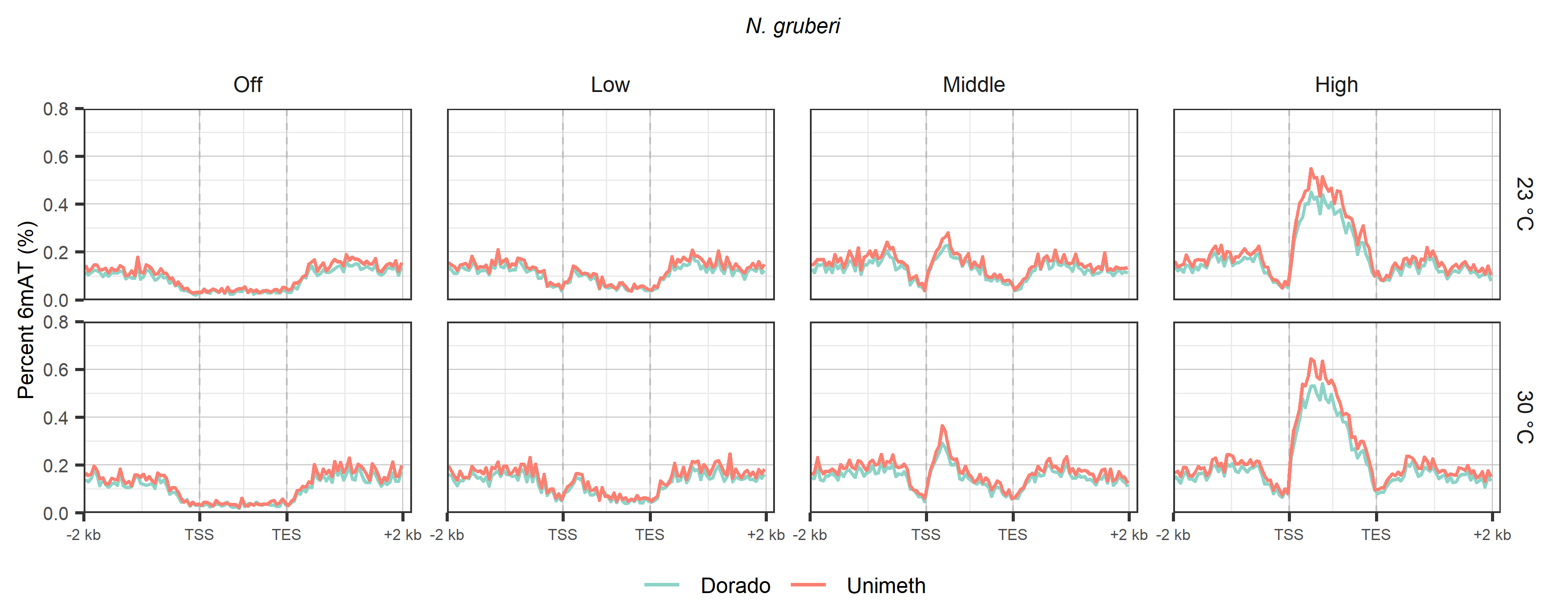

**Supplementary Fig. 28** Metaplots of 6mAT profiles across scaled gene regions in *N. gruberi* 23 °C and 30 °C samples, grouped by gene expression level (Off, Low, Middle, and High). For each group, 6mAT signal was averaged across a 1.5-kb scaled gene body with 2-kb upstream and downstream flanking regions, using 50-bp bins. Dashed vertical lines mark the transcription start site (TSS) and transcription end site (TES).

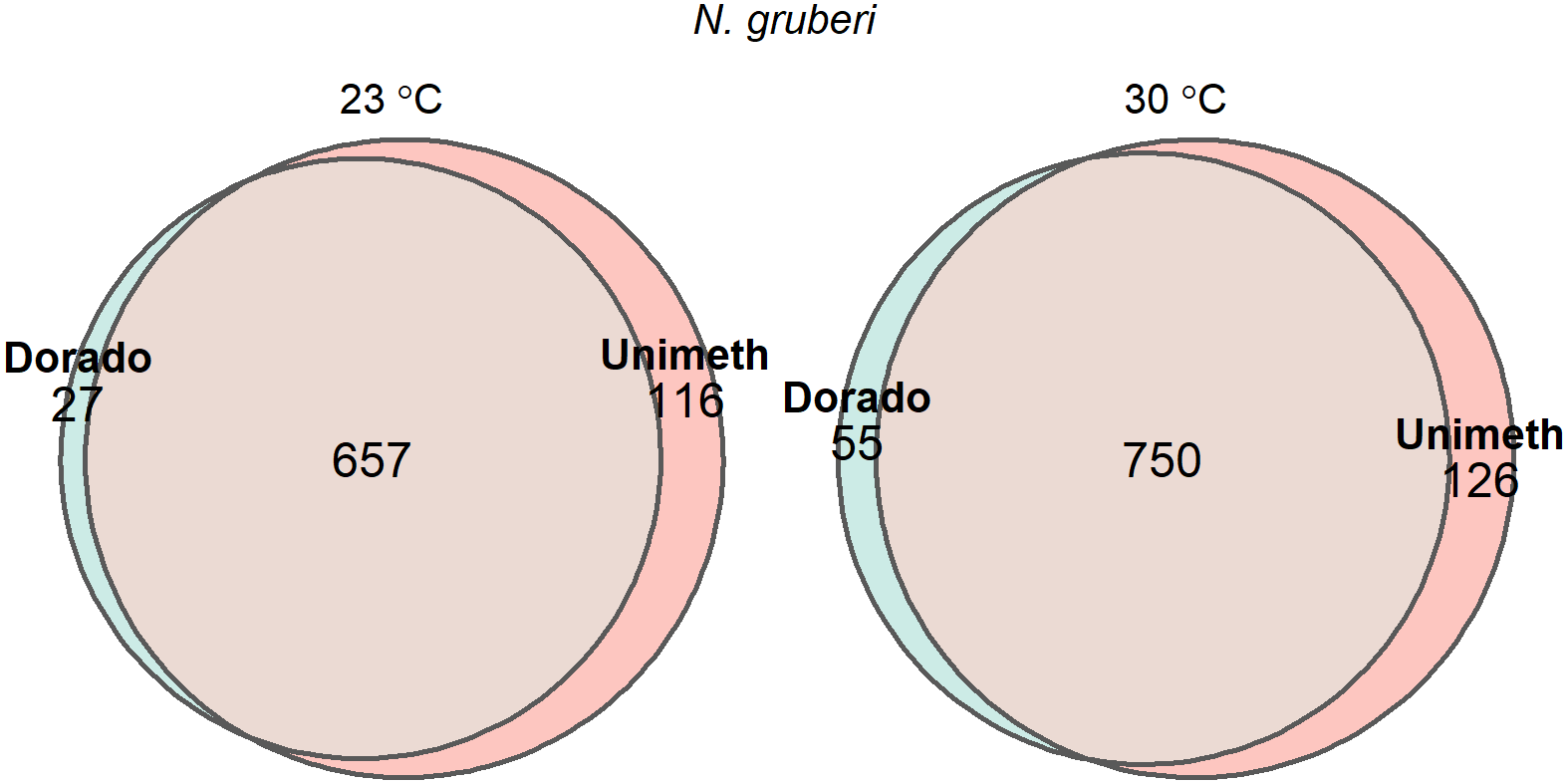

**Supplementary Fig. 29** Comparison of the numbers of 6mA-methylated genes identified by Dorado and Unimeth in *N. gruberi* samples grown at 23 °C and 30 °C.

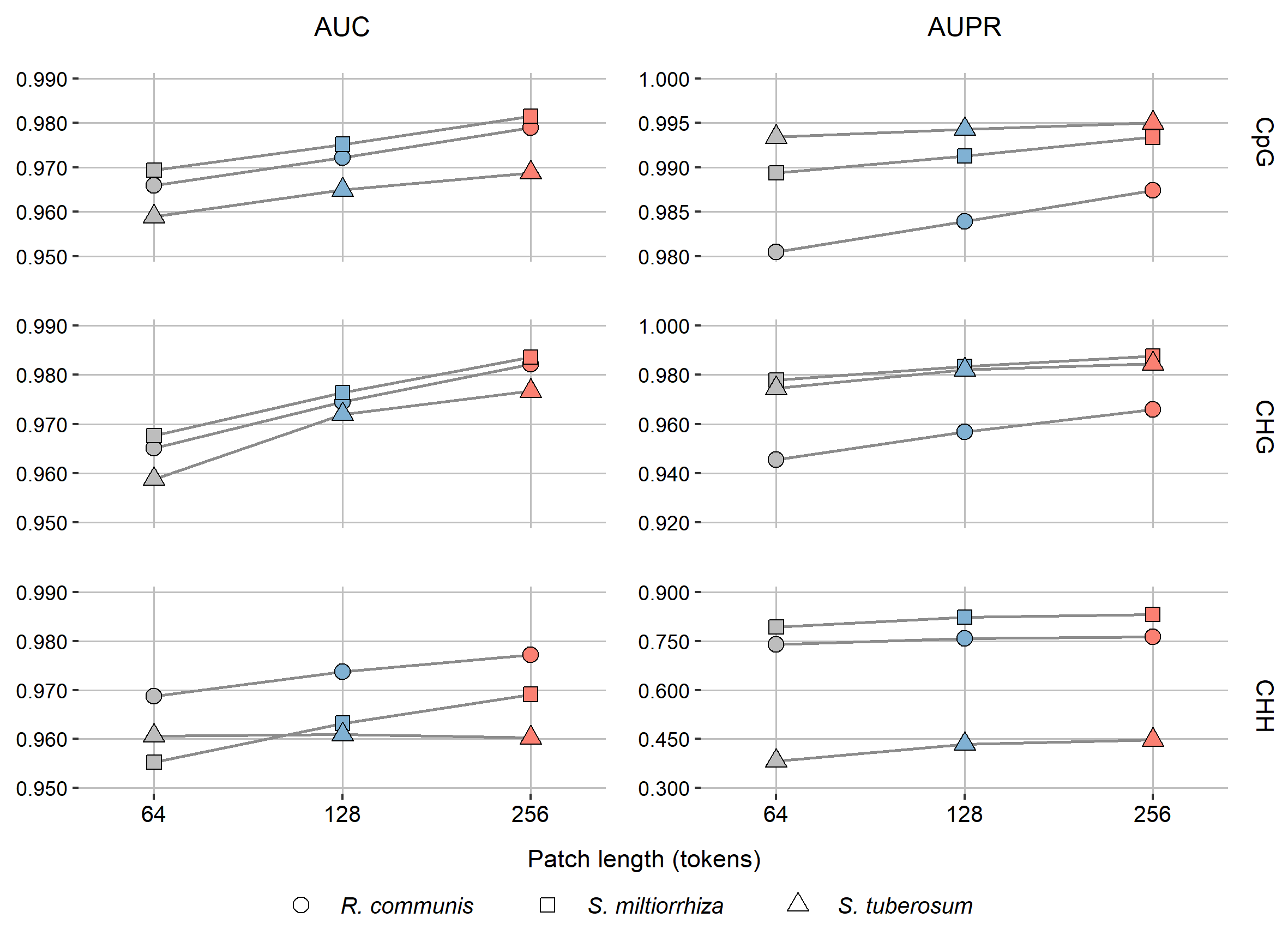

**Supplementary Fig. 30** Effect of patch length on Unimeth model performance. Models with patch lengths of 64, 128 and 256 tokens were fine-tuned from random initialization and evaluated across CpG, CHG and CHH contexts in R10.4.1 5kHz datasets from three plant species.

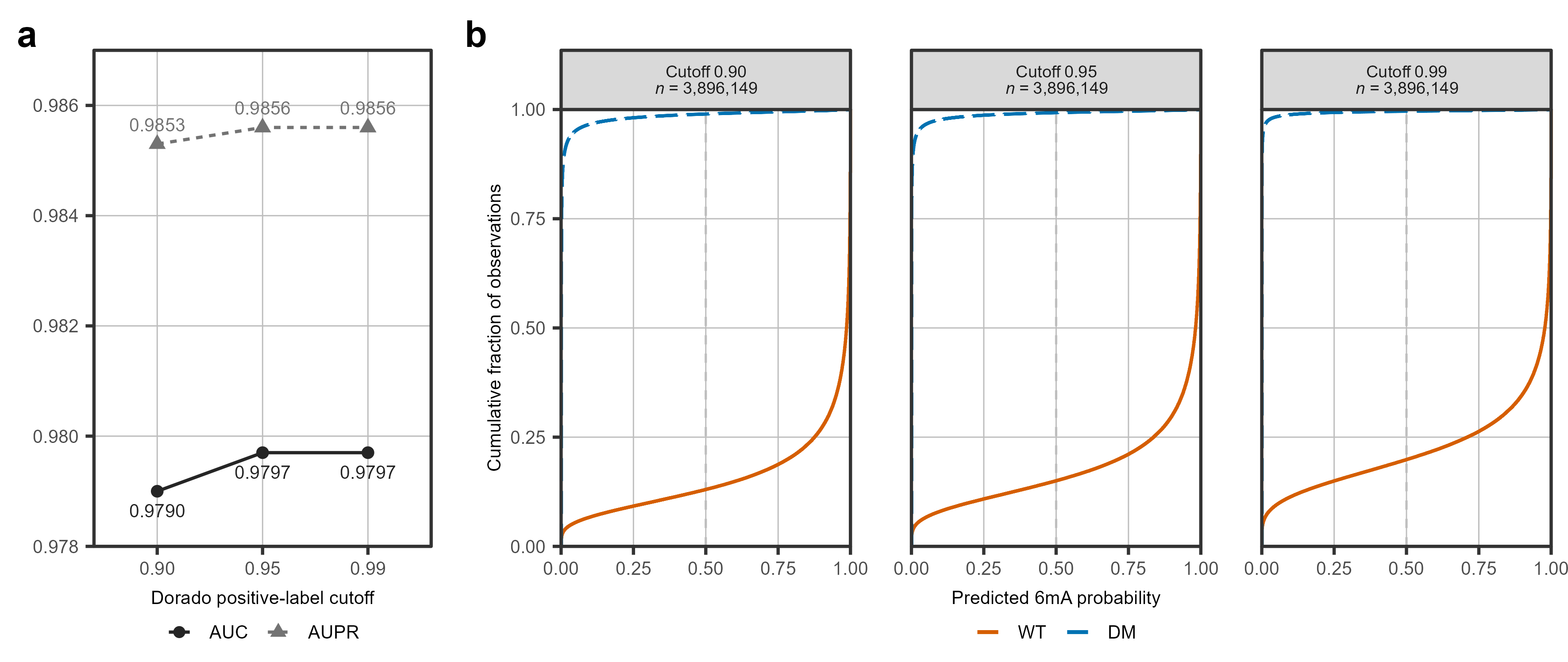

**Supplementary Fig. 31** Robustness of Unimeth 6mA detection to the Dorado probability cutoff used for positive pseudo-label selection. **a** AUC and AUPR of three Unimeth models trained using Dorado positive-label cutoffs of 0.90, 0.95 and 0.99 and evaluated on the *E. coli* wild-type (WT) versus *dam*/*dcm* methylase-knockout (DM) benchmark. **b** Empirical cumulative distributions of predicted 6mA probabilities at read-level GATC sites from WT and DM samples. Vertical dashed lines indicate the probability threshold of 0.5 used for binary methylation calling. *n* denotes the total number of GATC sites evaluated per model. Across the three models, AUC and AUPR remained stable, whereas increasing the positive-label cutoff shifted WT predictions towards lower probabilities.

### Supplementary Notes

**Supplementary Note 1 Reference genomes and annotations used in this study**

In this study, we analyzed sequencing data from 14 organisms, including 10 plants, 2 mammals, 1 heterolobosean protist, and 1 bacterium. The reference genomes and annotations used in this study are summarized below.

For *A. thaliana*, we downloaded the Col-CEN^2^ reference genome and annotations from <https://github.com/schatzlab/Col-CEN>. For *O. sativa*, we downloaded the T2T-NIP^3^ (AGIS1.0) assembly and corresponding annotations from <http://www.ricesuperpir.com/web/nip>.

For *Musa* spp., we generated a reference assembly using hifiasm^4^ (v0.25.0-r726) and purge_dups^5^ (v1.2.6). The hifiasm commands used for assembling the *Musa* spp. genome were:

*hifiasm -o file_prefix --ont -t 40 /path/to/Musa_spp/ont_reads/fastq*

*awk '/^S/{print ">"$2; print $3}' file_prefix.bp.p_ctg.gfa > file_prefix.bp.p_ctg.fa*

The resulting assembly was further processed with minimap2^6^ (v2.26-r1175) and purge_dups as follows:

*minimap2 -x map-ont -L -y -t 40 file_prefix.bp.p_ctg.fa /path/to/Musa_spp/ont_reads/fastq | gzip -c - > file_prefix.bp.p_ctg.align.paf.gz*

*pbcstat file_prefix.bp.p_ctg.align.paf.gz*

*calcuts PB.stat >cutoffs 2>calcults.log*

*python3 /path/to/purge_dups/scripts/hist_plot.py -c cutoffs PB.stat PB.cov.png*

*calcuts -l 23 -m 115 -u 220 PB.stat > cutoffs_manual*

*split_fa file_prefix.bp.p_ctg.fa > contigs.split.fasta*

*minimap2 -xasm5 -DP contigs.split.fasta contigs.split.fasta | gzip -c - > ctg2ctg.paf.gz*

*purge_dups -2 -T cutoffs_manual -c PB.base.cov ctg2ctg.paf.gz > dups.bed 2>purge_dups.log*

*get_seqs dups.bed file_prefix.bp.p_ctg.fa*

*cp purged.fa file_prefix.bp.p_ctg.fa.bp.p_ctg.purged.fa*

The file *file_prefix.bp.p_ctg.fa.bp.p_ctg.purged.fa* was used as the final reference genome. We used contig ptg000001l for method evaluation.

For the other 7 plant species, we used the same reference genomes as Chen *et al.*^1^. Reference genomes of 4 of these species were downloaded from NCBI, including *R. communis* ([GCF_019578655.1](https://www.ncbi.nlm.nih.gov/datasets/genome/GCF_019578655.1)), *S. lycopersicum* ([GCA_915070445.1](https://www.ncbi.nlm.nih.gov/datasets/genome/GCA_915070445.1)), *C. sinensis* ([GCF_022201045.2](https://www.ncbi.nlm.nih.gov/datasets/genome/GCF_022201045.2)), and *B. vulgaris* ([GCF_026745355.1](https://www.ncbi.nlm.nih.gov/datasets/genome/GCF_026745355.1)). For *S. miltiorrhiza*, *S. tuberosum*, and *V. vinifera*, we used the assembly results provided by Chen *et al.*^1^ as the reference genomes. For method evaluation, we used the longest contigs of these three assemblies, namely h2tg000013l, ptg000003l, and ptg000084l, respectively.

For the publicly available human datasets, we downloaded the CHM13 v2.0^7^ reference genome and gene annotation from the GitHub repository [marbl/CHM13](https://github.com/marbl/CHM13). We also downloaded annotations of peri/centromeric satellites, CpG islands, and repetitive genomic elements (RepeatMasker) from the corresponding UCSC Genome Browser tracks for T2T CHM13v2.0/hs1. For HBA20-2, we downloaded the GRCh38 primary assembly from GENCODE^8^ as the reference genome. For *M. musculus*, we downloaded the GRCm39 genome from GENCODE^8^ (<https://www.gencodegenes.org/mouse/release_M37.html>). For *E. coli*, we downloaded the reference genome [GCF_000005845.2](https://www.ncbi.nlm.nih.gov/datasets/genome/GCF_000005845.2/) from NCBI.

For *N. gruberi*, we used the re-assembled genome, corresponding gene annotation, and mRNA transcript sequences reported by Charria *et al.*^9^. These resources were used for analyses of 6mA distribution across gene regions and for integrating gene-level 6mA signals with RNA-seq data.

**Supplementary Note 2 Re-squiggle of R9.4.1 nanopore reads**

We employed the following strategy to re-squiggle the raw electrical signal values of R9.4.1 nanopore reads to the contiguous bases in the reference genome. First, reads were aligned to the reference genome using minimap2^6^ during basecalling. To maximize sequence continuity, unaligned terminal sequences were concatenated to their corresponding reference regions. Then, we processed the read-to-reference alignments by parsing the CIGAR strings generated by minimap2. We focused on handling deletions and insertions. For insertions, we appended raw signals to the signal of the last base in the preceding alignment segment. For deletions, we assigned empty signals to the corresponding base. Following re-squiggling, raw signals were normalized using the POD5 calibration parameters and the per-read signal-scaling parameters recorded by Dorado. We obtained *offset_POD5_* and *scale_POD5_* from the ‘calibration.offset’ and ‘calibration.scale’ fields, respectively, and *sm* and *sd* from the corresponding BAM tags. The ADC-space shift and scale were calculated by combining the POD5 ADC-to-pA calibration with the Dorado pA-to-normalized scaling (Equations (1) and (2)). The normalized signals were then calculated using Equation (3).

${shift}_{ADC}=\frac{sm}{{scale}_{POD5}}-{offset}_{POD5}$	(1)

${scale}_{ADC}=\frac{sd}{{scale}_{POD5}}$	(2)

${signal}_{norm}= \frac{{signal}_{raw} -{shift}_{ADC}}{{scale}_{ADC}}$	(3)

**Supplementary Note 3 Training implementation and hyperparameters**

Unimeth models were trained on eight NVIDIA A6000 GPUs using bfloat16 mixed precision. Model parameters were optimized using AdamW (β₁ = 0.9, β₂ = 0.999). Pre-training was run for 300,000 steps with a batch size of 384, a learning rate of 1×10⁻⁴ and a warm-up ratio of 0.05. Fine-tuning used a batch size of 288 and a learning rate of 1×10⁻⁵ without warm-up. The 5mC models were fine-tuned for 500,000 steps, and the 6mA model for 300,000 steps. When quantitative site-level references were available, calibration started from the corresponding fine-tuned checkpoint and was run for 10,000 steps with a batch size of 256 and a learning rate of 1×10⁻⁵.

**Supplementary Note 4 Detailed methods for 6mA evaluation and downstream analyses**

For 6mA model training, we used HG002 ONT Fiber-seq data generated with the R10.4.1 5kHz chemistry, excluding chromosome 1 for testing and downstream analyses. Dorado predictions were used to define training labels, with sites of 6mA probability ≥ 0.95 treated as positives and sites with probability = 0 in nucleosome regions predicted by *fibertools*^10^ treated as negatives. HG002 chromosome 1 Fiber-seq and matched native gDNA datasets were used for downstream evaluation of 6mA predictions. To compare global 6mA levels between methods, we calculated the fractions of methylated adenines across a range of methylation-calling thresholds for Unimeth and Dorado. For nucleosome analysis, nucleosome lengths were inferred from ONT 6mA calls using *fibertools* and compared with nucleosome profiles from PacBio Fiber-seq data. For nucleosome inference from ONT data, only predicted 6mA calls with probabilities ≥ 0.95 were retained. 10,000 reads were sampled for the analysis shown in Fig. 6b.

For independent evaluation of 6mA detection, we used an *E. coli* dataset containing wild-type (WT) and *dam/dcm* methylase-knockout mutant (DM) samples^11^. Read-level predictions were evaluated at GATC sites, with sites from WT reads treated as positive samples and sites from DM reads treated as negative samples. Model performance was assessed using the area under the receiver operating characteristic curve (AUC) and the area under the precision-recall curve (AUPR).

For *N. gruberi*, RNA-seq data from four samples, including two replicates each at 23 °C and 30 °C, were processed using fastp (v0.20.0), HISAT2 (v2.2.1), samtools (v1.21), StringTie (v2.1.2), and DeepTools (v3.5.5). RNA-seq reads were aligned to the *N. gruberi* reference genome using HISAT2 with “--rna-strandness RF” and a maximum intron length of 40,000 bp. Transcript abundance was quantified using StringTie with the “--rf” option, and gene expression levels were represented as TPM values. TPM values were averaged across replicates within each temperature condition for downstream analyses.

For *N. gruberi* 6mA analyses, we followed Charria *et al.*^9^ and used a probability threshold of 0.995 to define methylated 6mA sites in reads. Downstream analyses focused on ApT sites. Metagene-scale 6mA profiles were generated for annotated genes using regions spanning 2 kb upstream, scaled gene bodies, and 2 kb downstream. Only genes with annotated lengths ≥ 200 bp were retained, and gene bodies were scaled to 1,500 bp using 50-bp bins. For metagene-scale analyses, ApT sites with coverage ≥ 10 were retained and converted to methylation-percentage tracks for downstream profiling. For expression-stratified analyses, genes with TPM < 1 in all RNA-seq replicates were assigned to the off group, and the remaining genes were divided into low, middle and high expression groups according to tertiles of mean log2(TPM + 1). For gene-level 6mA analyses, methylated ApT sites were defined as sites with coverage ≥ 10 and methylation frequency > 10%. A gene was defined as methylated if it contained at least three such methylated ApT sites and had mean ApT coverage ≥ 10. Strand-paired ApT sites were collapsed into one locus. Based on gene-level 6mA calls, genes were classified as shared, Unimeth-only, Dorado-only, or unmethylated. Gene expression differences between methylation classes were assessed using the Wilcoxon rank-sum test.
